# Covalent Disruptor of YAP-TEAD Association Suppresses Defective Hippo Signaling

**DOI:** 10.1101/2022.05.10.491316

**Authors:** Mengyang Fan, Wenchao Lu, Jianwei Che, Nicholas Kwiatkowski, Yang Gao, Hyuk-Soo Seo, Scott B. Ficarro, Prafulla C. Gokhale, Yao Liu, Ezekiel A. Geffken, Jimit Lakhani, Kijun Song, Miljan Kuljanin, Wenzhi Ji, Jie Jiang, Zhixiang He, Jason Tse, Andrew S. Boghossian, Matthew G. Rees, Melissa M. Ronan, Jennifer A. Roth, Joseph D. Mancias, Jarrod A. Marto, Sirano Dhe-Paganon, Tinghu Zhang, Nathanael S. Gray

**Author notes:** Correspondence should be addressed to T.H.Z. and N.S.G. These authors contributed equally to this work (Mengyang Fan, Wenchao Lu).

## Abstract

The transcription factor TEAD, together with its coactivator YAP/TAZ, is a key transcriptional modulator of the Hippo pathway. Activation of TEAD transcription by YAP has been implicated in a number of malignancies, and this complex represents a promising target for drug discovery. However, both YAP and its extensive binding interfaces to TEAD have been difficult to address using small molecules, mainly due to a lack of druggable pockets. TEAD is post-translationally modified by palmitoylation that targets a conserved cysteine at a central pocket, which provides an opportunity to develop cysteine-directed covalent small molecules for TEAD inhibition. Here, we employed covalent fragment screening approach followed by structure-based design to develop an irreversible TEAD inhibitor MYF-03-69. Using a range of *in vitro* and cell-based assays we demonstrated that through a covalent binding with TEAD palmitate pocket, MYF-03-69 disrupts YAP-TEAD association, suppresses TEAD transcriptional activity and inhibits cell growth of Hippo signaling defective malignant pleural mesothelioma (MPM). Further, a cell viability screening with a panel of 903 cancer cell lines indicated a high correlation between TEAD-YAP dependency and the sensitivity to MYF-03-69. Transcription profiling identified the upregulation of proapoptotic *BMF* gene in cancer cells that are sensitive to TEAD inhibition. Further optimization of MYF-03-69 led to an *in vivo* compatible compound MYF-03-176, which shows strong antitumor efficacy in MPM mouse xenograft model via oral administration. Taken together, we disclosed a story of the development of covalent TEAD inhibitors and its high therapeutic potential for clinic treatment for the cancers that are driven by TEAD-YAP alteration.

## Introduction

The Hippo pathway is a highly conserved signaling pathway that regulates embryonic development, organ size, cell proliferation, tissue regeneration, homeostasis and is responsible for the development and progression of many malignancies^1^. Critical components of Hippo signaling pathway are TEAD transcription factors, that are present as a family of four highly homologous members (TEAD1-4) in mammals. Importantly, TEADs alone show minimal transcriptional activity and require binding with a coactivator YAP/TAZ to initiate efficient gene expression^2^. YAP has been well characterized as an oncoprotein^3^ and its tumorigenesis role is mostly associated with TEAD mediated YAP-dependent gene expression^4^. Tissue-specific deletion of the upstream regulators or overexpression of YAP itself results in hyperplasia, organ overgrowth and tumorigenesis^5^. Tumor suppressor Merlin, encoded by *NF2* gene, and kinase LATS1/2 are known upstream regulators that cooperatively promote phosphorylation of YAP and hence induce its retention and degradation in the cytoplasm^6^. However, loss of function mutations in *NF2* or *LATS1/2*, which occurs in >70% mesothelioma, promote YAP nuclear entry and binding with TEAD to drive oncogenesis^7^. In addition to mesothelioma, high level of nuclear YAP has been associated with poor prognosis in non-small cell lung cancer (NSCLC), pancreatic cancer, and colorectal cancer (CRC)^3a^. Moreover, there is evidence to suggest that activation of the YAP-TEAD transcriptional program can be involved with drug resistance^8^. For example, *ALK*-rearranged lung cancer cells were shown to survive the treatment of ALK inhibitor alectinib through YAP1 activation^9^. In EGFR-mutant lung cancers, YAP-TEAD was observed to promote cell survival and induce a dormant state upon EGFR signaling blockade^10^. Taken together, targeting YAP-TEAD has emerged as a promising therapeutic strategy for cancer treatment.

Over the decades, efforts to directly target YAP/TAZ-TEAD have focused on either designing peptide-mimicking agents that bind in the interface between YAP and TEAD and disrupt this interaction or using phenotypic screening to identify small molecules that inhibit YAP-dependent signaling^11^. For example, a cyclic YAP-like peptide has been developed as a potent disruptor of YAP-TEAD complex in cell lysates but failed to exert effects in cells due to poor cellular permeability^12^. Recently, TEAD stability and the association with YAP protein was shown to be regulated by *S*-palmitoylation, a post-translational modification on a conserved cysteine located within the palmitate binding pocket (PBP) on YAP binding domain (YBD) of TEAD^13^. This finding led to the discovery of several small molecules: flufenamic acid (FA)^14^, MGH-CP1^15^, compound 2^16^ and VT101∼107^17^, that reversibly bind TEAD PBP (**Figure S1**). Additionally, inspired by thioester formation of the conserved cysteine with palmitate, covalent TEAD inhibitors TED-347^18^, DC-TEADin02^19^ and K-975^20^ (**Figure S1**) have also been designed. TED-347 and DC-TEADin02 covalently bind with TEAD using a highly electrophilic chloromethyl ketone and vinyl sulfonamide warhead. In comparison, a less electrophilic arylamide warhead is used in the design of K-975, which shows a potent cellular and *in vivo* activity in mouse tumor models albeit employing a high dose. Although these compounds have adequately demonstrated that the palmitoylated cysteine is targetable, none of them are optimized compounds and the structure-activity study for each of these scaffolds has not been disclosed. We have previously reported MYF-01-37 which possesses a less electrophilic warhead compared to K-975 as a covalent TEAD inhibitor and demonstrated its utility to blunt the transcriptional adaptation in mutant EGFR-dependent NSCLC cells^10^. However, MYF-01-37 is a sub-optimal chemical probe which requires micromolar concentrations in cellular assays and displays poor pharmacokinetic properties.

Development of covalent chemical probes and drugs has experienced a revival in recent years resulting in a deluge of new modality targeting a range of cancer-relevant targets, such as KRAS^G12C^, EGFR, BTK, JAK3 and others^21^. The conventional approach to develop a covalent inhibitor is using a structure-guided design to equip a pre-existing reversible binder with an electrophile to target a nucleophilic residue on the target protein, most typically a cysteine. Alternatively, a recently developed approach involves screening a library of low molecular weight electrophilic fragments followed by medicinal chemical optimization^22^. Here, we used such screening approach to identify the lead covalent fragment MYF-01-37. Subsequent structure-based design to engage a side pocket resulted in development of a more potent covalent TEAD inhibitor MYF-03-69, which inhibits the palmitoylation process of all four TEAD paralogs *in vitro*, disassociates the endogenous YAP-TEAD complex, downregulates YAP-dependent target gene expression, and preferentially impairs the proliferation of mesothelioma cells that exhibit defects in Hippo signaling. A comprehensive screen against 903 cancer cell lines^23^ identified more YAP-TEAD dependent cells as sensitive to MYF-03-69 in addition to mesothelioma. Transcription profiling suggests that the upregulation of Bcl-2 modifying factor (*BMF)* gene correlated with antiproliferative response to TEAD inhibition. Further, *in vivo* pharmacological effect was achieved with orally bioavailable compound MYF-03-176, indicating a promising lead for therapy of cancers driven by Hippo signaling dysfunction.

## Results

### Covalent fragment screening identified MYF-01-37 as a covalent TEAD binder

To screen for a covalent lead compound for TEAD, we first assembled a biased covalent fragment library that was developed based on the flufenamic acid template and a cysteine-reactive acrylamide warhead^24^ (**Figure 1a**, **Figure S2a**). We then conducted a medium-throughput screen of this library using mass spectrometry (MS) to detect protein-fragment adduct formation with recombinant, purified TEAD2 YBD protein. MYF-01-37 was identified as the fragment capable of efficient labeling while the majority of the fragment library failed to label the protein (**Figure S2b**). Next, the protein labeled with MYF-01-37 was proteolytically digested and the resultant peptides were analyzed using tandem mass spectrometry (MS/MS) which confirmed labeling of C380, which is the cysteine previously reported to be *S*-palmitoylated in TEAD2^13a^ (**Figure 1a**).

**Figure 1.**
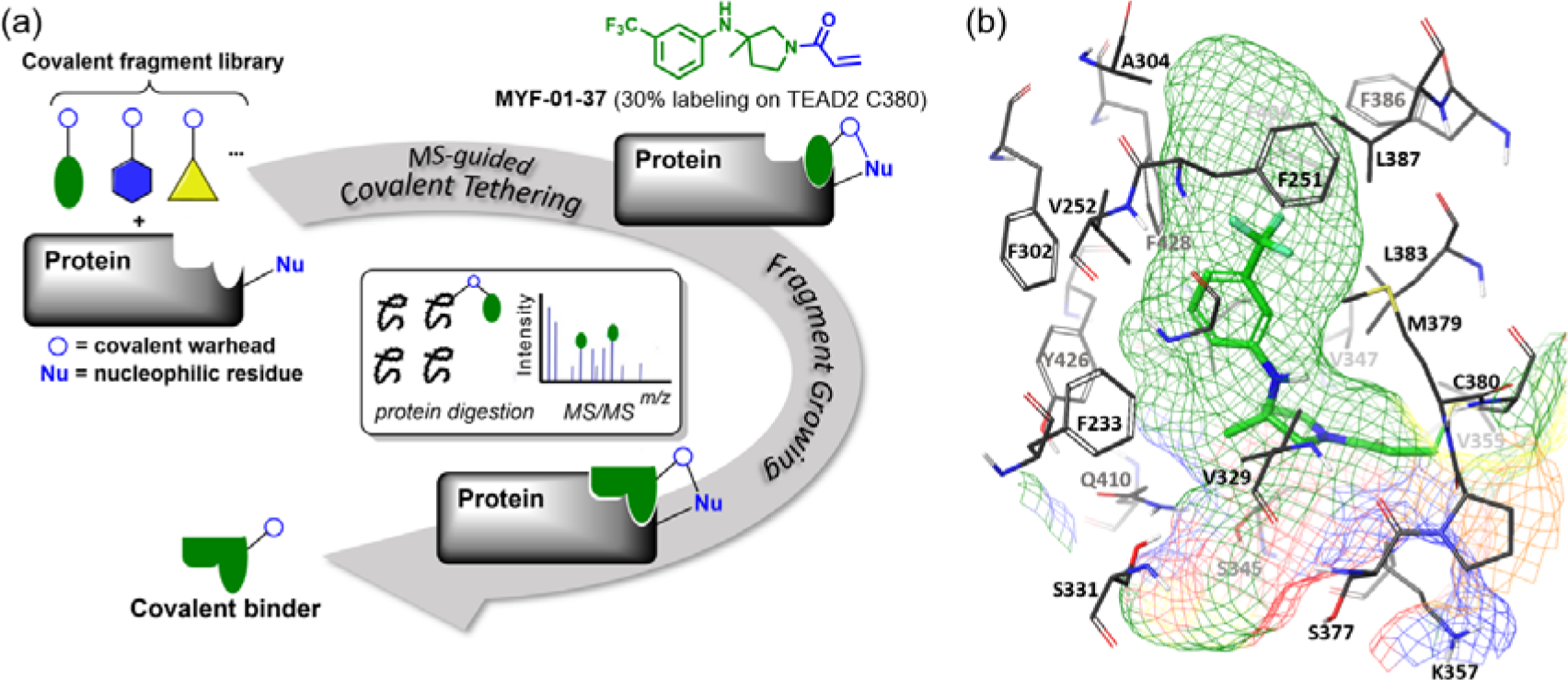
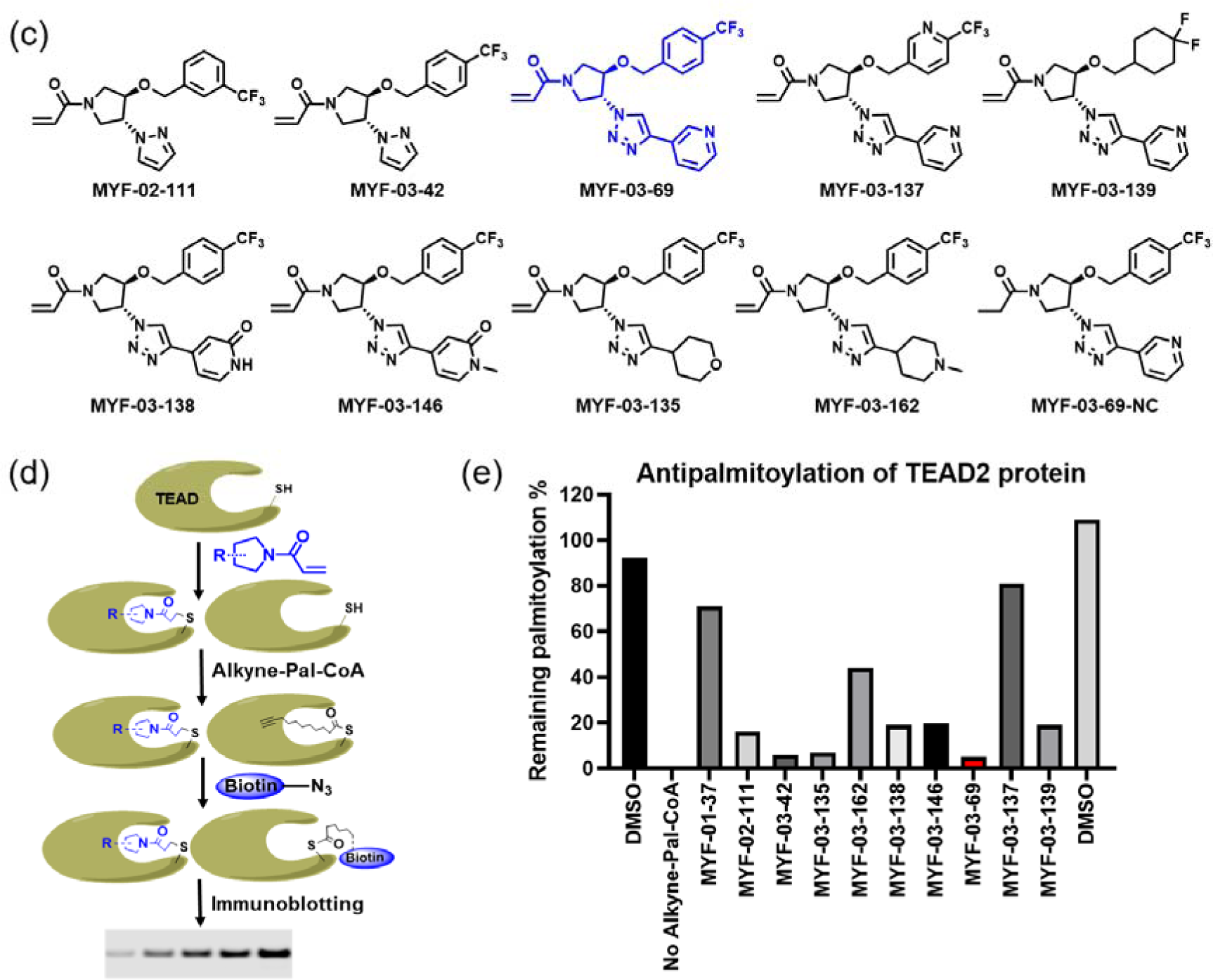
Covalent fragments screening and structure-guided design to identify Y-shaped compound MYF-03-69. (**a**) Illustration of covalent fragment library screening and optimization for TEAD inhibitor. (**b**) Surface of TEAD2 palmitate pocket depicted in mesh with MYF-01-37 (in green) modeled in. Residues forming the pocket were labeled. The color of mesh indicates hydrophobicity of the pocket surface. (**c**) Chemical structures of representative Y-shaped compounds. (**d**) The schematic diagram of *in vitro* palmitoylation assay. (**e**) Anti-palmitoylation activity of MYF-01-37 and Y-shaped compounds after TEAD2 protein was pre-incubated with 2 μM compound at 37°C for 2 hours.

### Structure-based optimization yields MYF-03-69, an irreversible inhibitor of TEAD palmitoylation

Next, we employed a structure-based fragment growing strategy to elaborate MYF-01-37 to develop a more potent and selective inhibitor (**Figure 1a**). Given that the site of covalent modification is located at the opening of the palmitate binding pocket, we analyzed the TEAD palmitate pockets from multiple crystal structures available in the Protein Data Bank (PDB)^13–14^. All the structures aligned well, indicating a conserved and relatively inflexible pocket. As shown in the **Figure 1b**, docking of MYF-01-37 and TEAD2 revealed that MYF-01-37 binds in a tunnel formed by side chains of hydrophobic residues F233, F251, V252, V329, V347, M379, L383, L387, F406 and F428. We also identified a side pocket lined with hydrophilic side chains contributed by S331, S345, S377 and Q410. We hypothesized that we could improve potency and selectivity for our covalent ligands by introducing polar interactions and shape complementarity to this side pocket. Therefore, we predicted that the optimized binding pocket is Y-shaped with the targeted cysteine at one end, and hydrophobic and hydrophilic pockets at the other two ends. We envisioned that occupying the hydrophilic side pocket will also provide chance to incorporate polar moiety, leading to reduced global hydrophobicity and more druglike molecules. With this rationale, a variety of hydrophilic groups were introduced to the pyrrolidine ring, leading to synthesis of a focused library of Y-shaped molecules, with representatives shown in **Figure 1c**. We employed an *in vitro* palmitoylation assay that uses an alkyne-tagged palmitoyl CoA as the lipidation substrate and clickable biotin (**Figure 1d**), to screen this series of compounds for their ability to inhibit palmitoylation (**Figure 1e**, **Figure S3**). We observed that the *para* substituted trifluoromethyl benzyl is superior to *meta* substituted trifluoromethyl benzyl (MYF-02-111 vs. MYF-03-42). We then replaced pyrazole by a substituted triazole to explore the hydrophilic pocket (MYF-03-135, MYF-03-162, MYF-03-138, MYF-03-146, MYF-03-69), and observed a sensitivity to various substitutions at C4 position of the triazole ring. For example, tetrahydropyran and pyridine enhanced the activities, whereas piperidine and pyridone were much less favored. In addition, we noticed that extending the hydrophobic tail of MYF-01-37 allowed more complete occupancy of the hydrophobic channel. Further modifications of the groups occupying the hydrophobic tunnel did not yield further improvement on the potency compared to trifluoromethyl benzyl (MYF-03-137, MYF-03-139), implicating restrict requirement for a hydrophobic moiety. Amongst this series of compounds, we identified MYF-03-69 (**Figure 1c**) as the most potent compound, and chose it for further characterization.

### MYF-03-69 occupies the Y-shaped pocket and forms covalent bond with the conserved cysteine of TEAD

Given that labeling efficiency of the parental fragment MYF-01-37 was moderate, we examined whether the elaborated, Y-shaped molecule labeled TEAD more efficiently. Recombinant TEAD2 YBD was incubated with MYF-03-69, followed by mass spectrometry. Incubation with 10-fold molar excess of MYF-03-69 at room temperature for 1 hour led to 100% labeling (**Figure 2a**) and follow-up trypsin digestion and MS/MS analysis confirmed C380 as the modified residue (**Figure 2b**). This covalent bond formation and the exact binding mode was further validated by obtaining a high-resolution co-crystal structure of TEAD1 YBD in complex with MYF-03-69 (1.68 Å) (**Figure 2c**). Unambiguous electron density of a covalent bond was observed between TEAD1 C359 (analogous to C380 in TEAD2) and the acrylamide warhead of MYF-03-69 with the remaining part of the molecule adopting a Y-shape and establishing specific interactions with the hydrophobic pocket and hydrophilic patch as predicted (**Figure 2c**). MYF-03-69 was completely buried in the lipid pocket with the *para*-trifluoromethyl benzyl group extended along a lipid trajectory, forming extensive hydrophobic interactions with side chains of A223, F239, V240, A292, I366, and F407. The 3-pyridinyl group occupied the side pocket, forming a favorable water-bridged hydrogen bond network with adjacent S310 and Y312 via pyridinyl nitrogen (**Figure 2d**). Overall, high resolution structural analysis confirms that MYF-03-69 forms covalent attachment with the TEAD cysteine that is the site of *S*-palmitoylation, and that the elaborated molecule forms specific interactions via both hydrophobic and hydrophilic regions surrounding the targeted cysteine.

**Figure 2.**
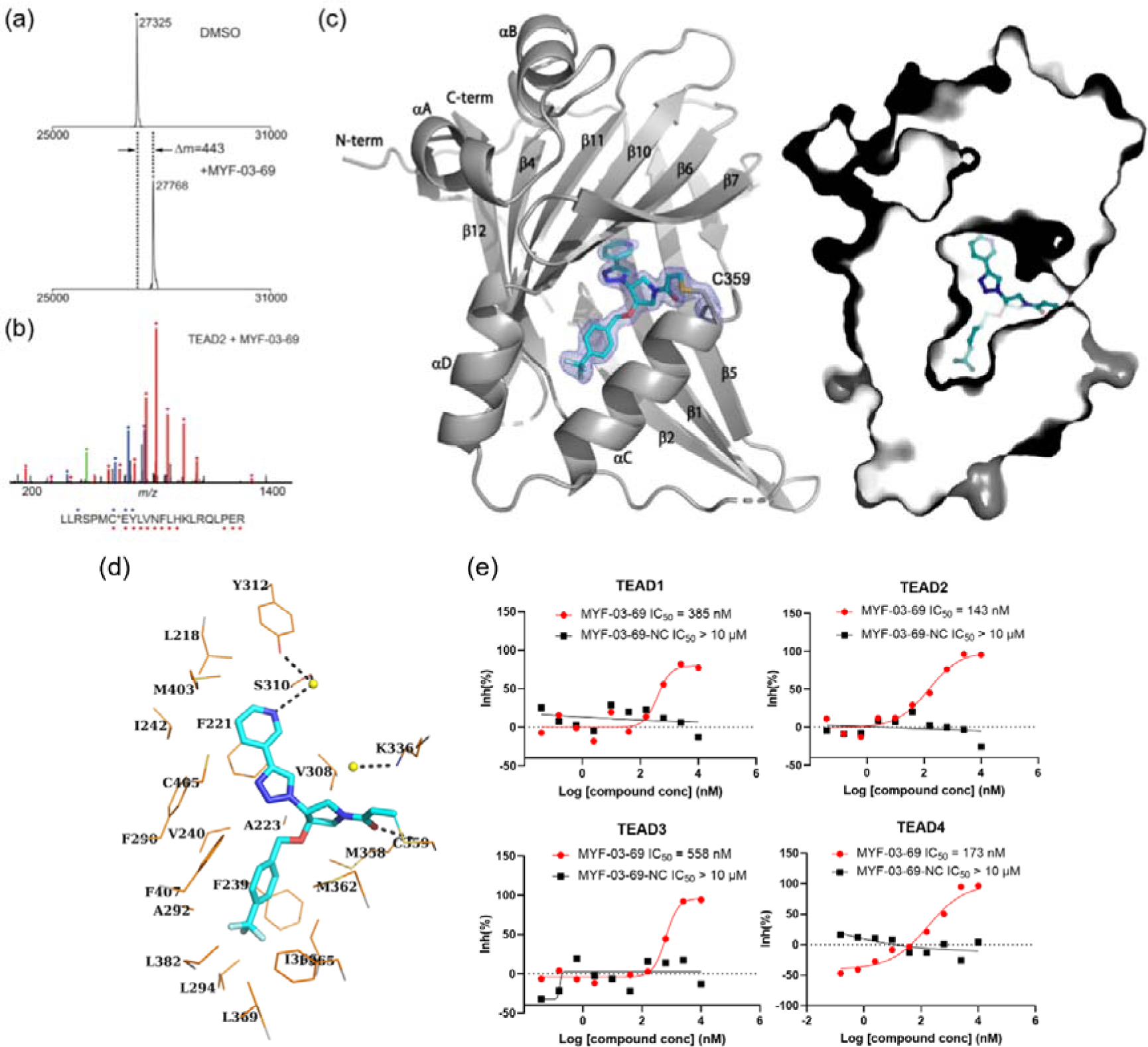
MYF-03-69 binds in TEAD palmitate pocket covalently through the conserved cysteine. (**a**) The mass labeling of intact TEAD2 protein by MYF-03-69. (**b**) Trypsin digestion and tandem mass spectrum (MS/MS) localize labeling site of cysteine 380. (**c**) Co-crystal structure of MYF-03-69 with TEAD1 indicates covalent bond formation with Cys359. The compound adopts Y-shape and binds in both lipid tunnel and hydrophilic side pocket. (**d**) Interactions between MYF-03-69 and TEAD1 palmitate pocket. (**e**) A dose titration of MYF-03-69 and MYF-03-69-NC in anti-palmitoylation assay on TEAD1-4. Recombinant YBD protein of TEADs were preincubated with compounds at 37°C for 2 hours. Data are representative of n = 3 independent experiments.

To further characterize biochemical activity of MYF-03-69, we performed a dose titration for all TEAD paralogs (**Figure 2e**, **Figure S4**). Preincubation of all recombinant TEAD1-4 YBDs with MYF-03-69 potently inhibited palmitoylation with similar IC_50_ values at submicromolar concentrations, indicating that MYF-03-69 is a pan-TEAD inhibitor. This is consistent with the high sequence conservation of residues in the palmitate pockets of TEAD1-4. In contrast, MYF-03-69-NC (**Figure 1c**), the non-covalent counterpart of MYF-03-69 incapable of forming covalent bond, completely lost activity across all TEADs, which demonstrated the essentiality of covalent bond formation.

### MYF-03-69 inhibits TEAD palmitoylation and disrupts endogenous YAP-TEAD association in cells

We next investigated the ability of MYF-03-69 to modulate TEAD palmitoylation inside cells with ω-alkynyl palmitic acid (Alkyne-C16) as probe. Alkyne-C16 contains an alkyne group at C15 and has been shown to be metabolically incorporated into cellular proteins at native palmitoylation sites. The previously reported pulse-chase style experiment^16^ was adopted and modified to monitor TEAD palmitoylation in HEK293T cells transfected with Myc-TEAD4. After the cells were incubated with Alkyne-C16 for 24 hours, the labeled Myc-TEAD4 was conjugated with biotin-azide through a Cu(I)-catalyzed click reaction. With western-blotting, we found that the alkyne-palmitate labeling of Myc-TEAD4 decreased after cotreatment with varying concentrations of MYF-03-69 for 24 hours compared to the DMSO treatment, while TEAD protein levels were not affected (**Figure 3a**). This result suggested that MYF-03-69 competed off palmitic acid during TEAD lipidation in cells. Consistently with biochemical result, the negative control compound MYF-03-69-NC did not exhibit any effect on TEAD (**Figure 3a**).

**Figure 3.**
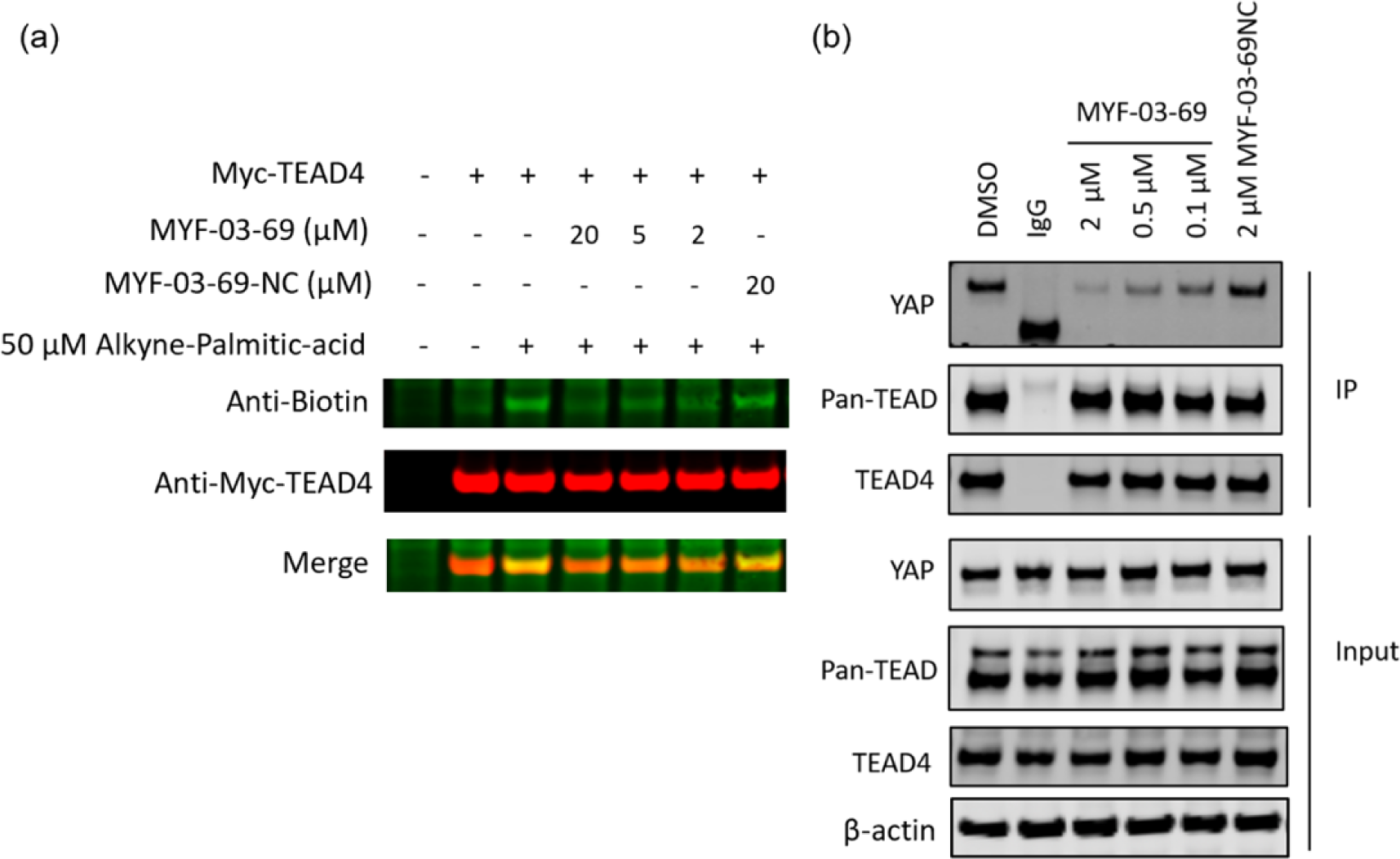
MYF-03-69 inhibits palmitoylation of TEAD protein and disrupts its association with YAP in cells. (**a**) Palmitoylation of Myc-TEAD4 in HEK293T cells after treatment with MYF-03-69 and MYF-03-69-NC indicated by an alkyne-palmitic-acid probe and click chemistry. Cells were treated for 24 hours. (**b**) Co-immunoprecipitation (Co-IP) of endogenous YAP and TEAD in NCI-H226 cells after treatment with MYF-03-69 and MYF-03-69-NC at indicated doses. Cells were treated for 24 hours.

Palmitoylation of TEADs was recently proposed to be required for their binding to YAP and TAZ^13b^. Although TEAD inhibitors with different chemotypes have been recently developed, it remains controversial whether pharmacological targeting of the TEAD palmitate pocket could disrupt YAP/TAZ association^25^. In order to investigate whether MYF-03-69 impairs YAP-TEAD interaction, we conducted an endogenous co-immunoprecipitation (co-IP) experiment to monitor YAP-TEAD interactions in the presence of the compound. Notably, 24-hour treatment of NCI-H226 cells with MYF-03-69 significantly decreased endogenous YAP co-immunoprecipitation with a pan-TEAD antibody, whereas this treatment had minimal effect on TEAD protein level (**Figure 3b**). In parallel, we also evaluated MYF-03-69-NC and observed no effect on YAP-TEAD association in cells. Taken together, these results suggest that MYF-03-69-mediated perturbation of TEAD palmitoylation disrupts YAP and TEAD protein interactions.

To demonstrate MYF-03-69 as a selective covalent TEAD binder, we interrogated the proteome-wide reactivity profile of MYF-03-69 on cysteines using a well-established streamlined cysteine activity-based protein profiling (SLC-ABPP) approach (**Figure S5a**)^26^. We employed the cysteine reactive desthiobiotin iodoacetamide (DBIA) probe^26^ which was reported to map more than 8,000 cysteines and performed a competition study in NCI-H226 cells pretreated with 0.5, 2, 10 or 25 µM of MYF-03-69 for 3 hours in triplicate. The cysteines that were conjugated >50% (competition ratio CR>2) compared to DMSO control were analyzed and assigned to the protein targets (**Figure S5b**). In the DMSO control group, although DBIA mapped 12,498 cysteines in total, the TEAD PBP cysteines were not detected. Among 12,498 mapped cysteines, only 7 cysteines were significantly labeled (i.e. exhibited >50% conjugation or CR>2) by 25 µM of MYF-03-69, and all of these sites exhibited dose-dependent engagement (**Supplementary Dataset 1**). The missing TEAD cysteines might be due to low TEAD1-4 protein abundance and/or inability of the PBP cysteines to be labeled by DBIA. DBIA probe can only be applied in cell lysate context, which might also result in the missing labeling on TEAD. To our knowledge, all the proteins with meaningful labeling efficiency (CR>2) cysteine sites are not known to be involved in the YAP/TAZ-TEAD signaling pathway. Taken together, our proteomic analysis suggests that MYF-03-69 exhibited quite low reactivity towards the cysteine proteome, although TEAD PBP cysteines were not detected by the DBIA probe.

### MYF-03-69 inhibits TEAD transcription and downregulates YAP target genes expression in mesothelioma cells

The majority of mesothelioma patients (50-75%) harbor genetic alterations in Hippo pathway regulatory components, including *NF2* loss of function mutation/deletion and *LATS1-PSEN1* fusion/*LATS2* deletion, which lead to YAP activation and Hippo pathway gene expression^7^. Thus, in order to monitor YAP-TEAD transcriptional activity, Hippo/YAP reporter cells were generated in NCI-H226, a *NF2*-deficient MPM cell line. After 72-hour treatment of MYF-03-69, YAP-TEAD transcriptional activity of reporter cells significantly decreased in a dose-dependent manner with an IC_50_ value of 56 nM, while MYF-03-69-NC had minimal effect (**Figure 4a**). The transcriptional inhibition also led to significant downregulation of canonical YAP target genes including *CTGF*, *CYR61*, *IGFBP3*, *KRT34* and *NPPB* and upregulation of proapoptotic gene *BMF* in NCI-H226 cells and MSTO-211H, a *LATS1*-*PSEN1* fusion MPM cell line (**Figure 4b**-**c**), while it showed much milder effects on transcription in normal mesothelium cells MeT-5A (**Figure S6**). Protein level of *CYR61* and *AXL* gene products also decreased (**Figure 4d**, **Figure S7**). Encouraged by these results, we set out to study whole transcriptome perturbation by MYF-03-69. RNA sequencing was performed in NCI-H226 cells that were treated with 0.1 μM, 0.5 μM, and 2 μM of MYF-03-69. There were 339 genes that exhibited significant differential expression at 2 µM treatment, and the majority of them changed in a dose dependent manner (**Figure 4e** and **Supplementary Dataset 2**). The genes that were differentially expressed with statistical significance (Fold change ≥ 1.5 and adjusted p value ≤ 0.05) at 2 μM treatment condition, are colored in red and partially labeled in the volcano plot (**Figure 4f**). Compared to high concentration, the number of genes with significantly altered expression level dropped off quickly at lower concentrations (**Figure S8a-b**). For example, at the concentration of 0.1 μM, only *CTGF* showed statistically significant change (**Figure S8b**). While multiple YAP target genes such as *CTGF*, *ADM*, *ANKRD1* were significantly downregulated^2^, *DDIT4* was observed to be upregulated^27^. To investigate whether the 339 differentially expressed genes were concentrated in particular biological pathways, KEGG pathway enrichment calculation^28^ were carried out, which demonstrated Hippo signaling pathway was among the top 5 enriched processes (**Figure 4g**). Beyond the Hippo pathway, gap junction^29^ and WNT signaling pathway^30^ were also enriched, consistent with pleiotropic functions and cross-talk of YAP-TEAD pathway in diverse biological processes. Overall, we documented that MYF-03-69 affects transcription in a manner consistent with its activity as a disruptor of YAP-TEAD interactions.

**Figure 4.**
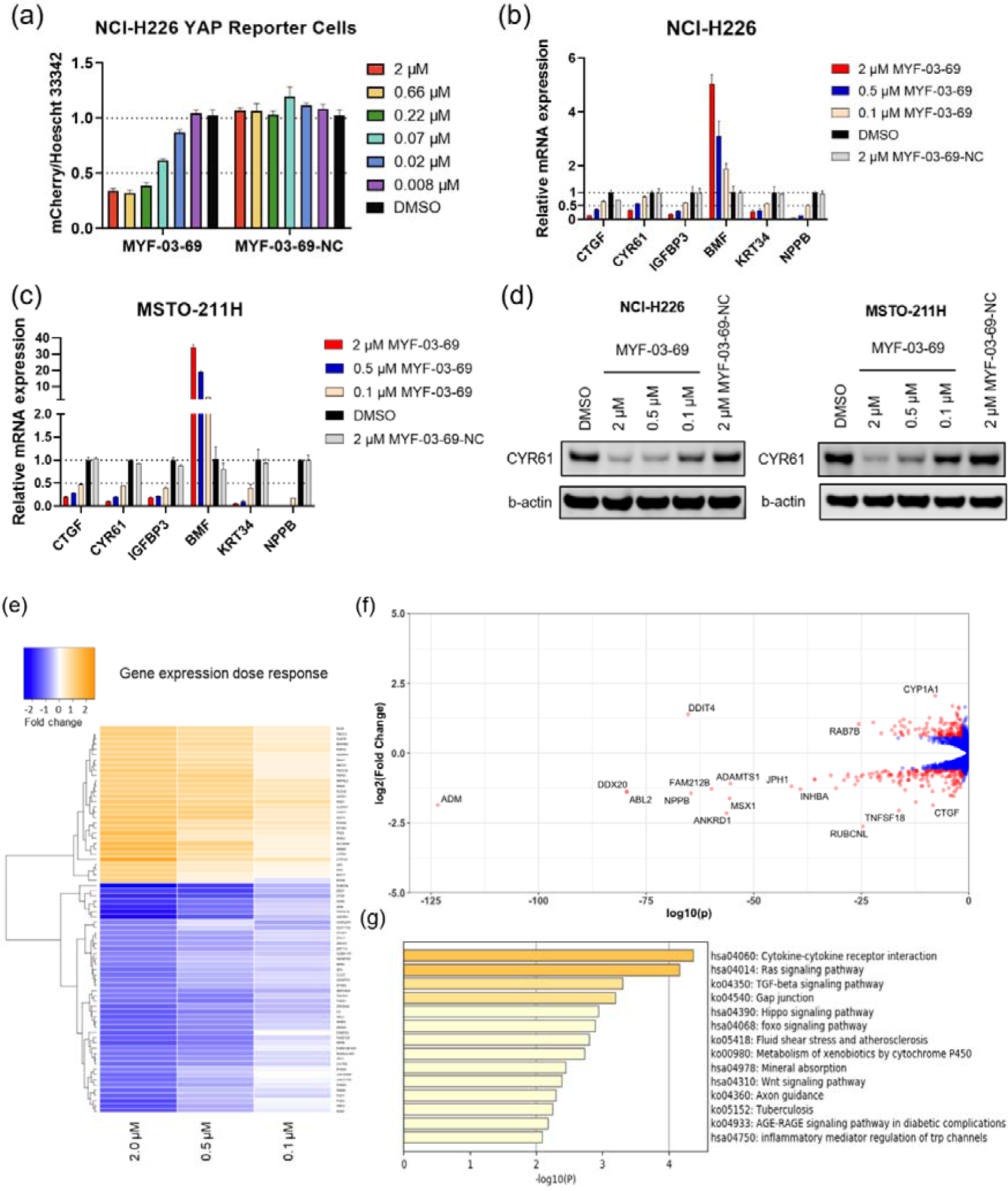
MYF-03-69 inhibits YAP-TEAD transcription. (**a**) MYF-03-69, but not MYF-03-69-NC, inhibits YAP-TEAD transcriptional activity in NCI-H226 mCherry reporter cells. Cells were treated for 72 hours. (**b**-**c**) MYF-03-69, but not MYF-03-69-NC, downregulates YAP target genes and upregulates a pro-apoptotic gene *BMF*. Cells were treated for 24 hours. Data were presented as mean ± SD of n = 3 biological independent samples. (**d**) MYF-03-69, but not MYF-03-69-NC, downregulates CYR61 protein level in NCI-H226 and MSTO-211H cells. (**e**) Heatmap for gene expression change with MYF-03-69 treatment at indicated concentrations. (**f**) Differential gene expression from RNA sequencing of NCI-H226 cells treated with 2 μM MYF-03-69. The differentially expressed genes with FC ≥ 1.5 and p ≤ 0.05 were colored in red and labeled. Not all differentially expressed genes were labeled. (**g**) Pathway enrichment analysis of differentially expressed genes from 2 μM compound treatment samples.

### MYF-03-69 selectively inhibits mesothelioma cancer cells with defective Hippo signaling

Next, we investigated anti-cancer activity of MYF-03-69 on Hippo signaling defective mesothelioma cells. As shown in **Figure 5a** and **Figure S9a**, 5-day cell growth assays demonstrated that MYF-03-69 potently retarded the cell growth of NCI-H226 and MSTO-211H, while it showed no antiproliferation activity against MeT-5A and NCI-H2452 cells, which are non-cancerous mesothelium cells and mesothelioma cells with intact Hippo signaling, respectively. These antiproliferative effects in NCI-H226 and MSTO-211H cells were observed under 3D spheroid suspensions culture condition as well (**Figure 5b**). Under the same conditions, the negative control compound MYF-03-69-NC was not antiproliferative (**Figure 5b**, **Figure S9b**). Further, cell cycle analysis demonstrated that 48-hour treatment with MYF-03-69 on NCI-H226 and MSTO-211H cells caused cell cycle arrest at the G1 phase, which is in accordance with previous findings obtained from genetic knockdown of YAP^31^, while negative control compound had no effect (**Figure 5c**-**d**). Collectively, these results show that inhibition of TEAD palmitoylation by MYF-03-69 effectively and precisely affects YAP-TEAD function.

**Figure 5.**
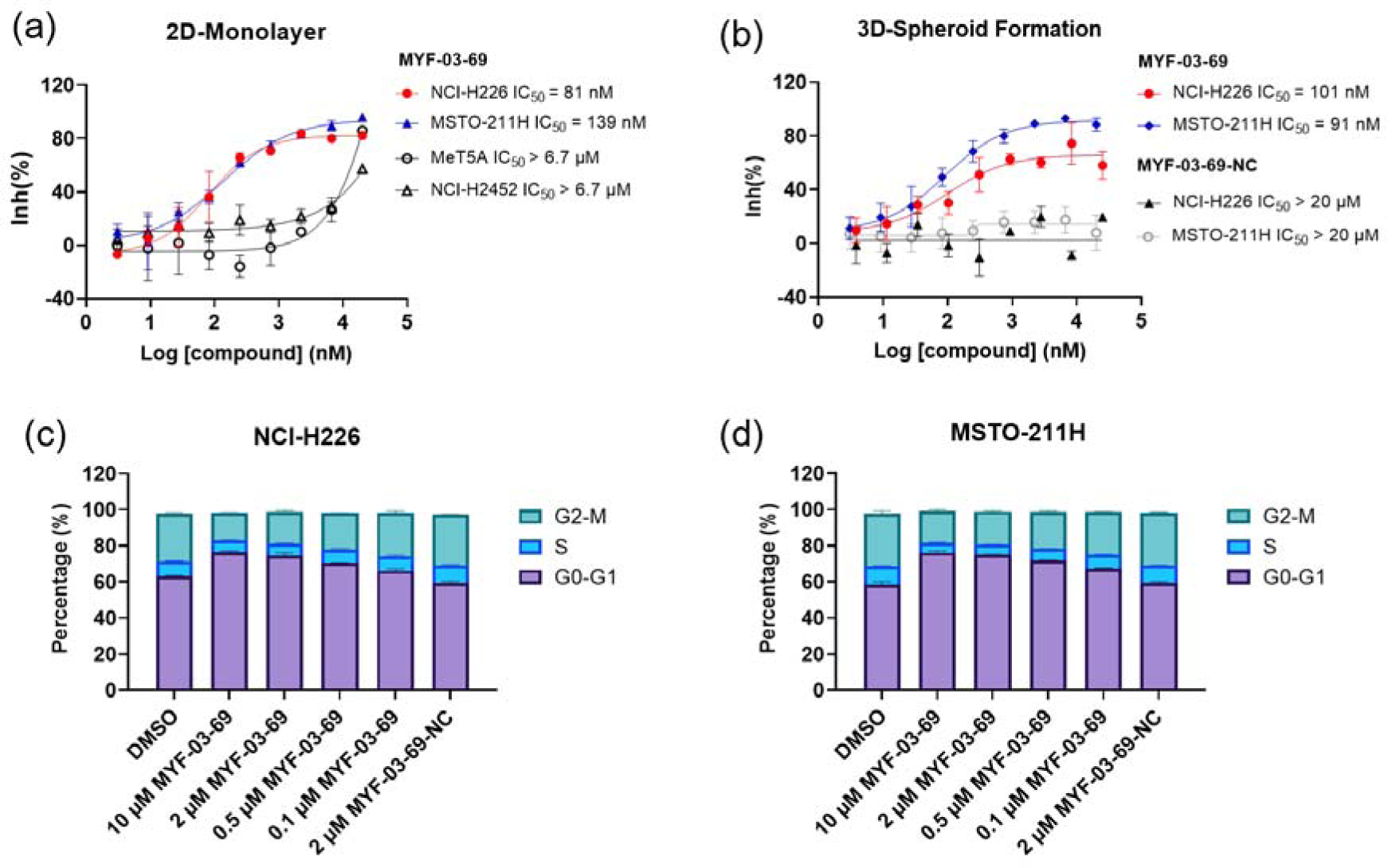
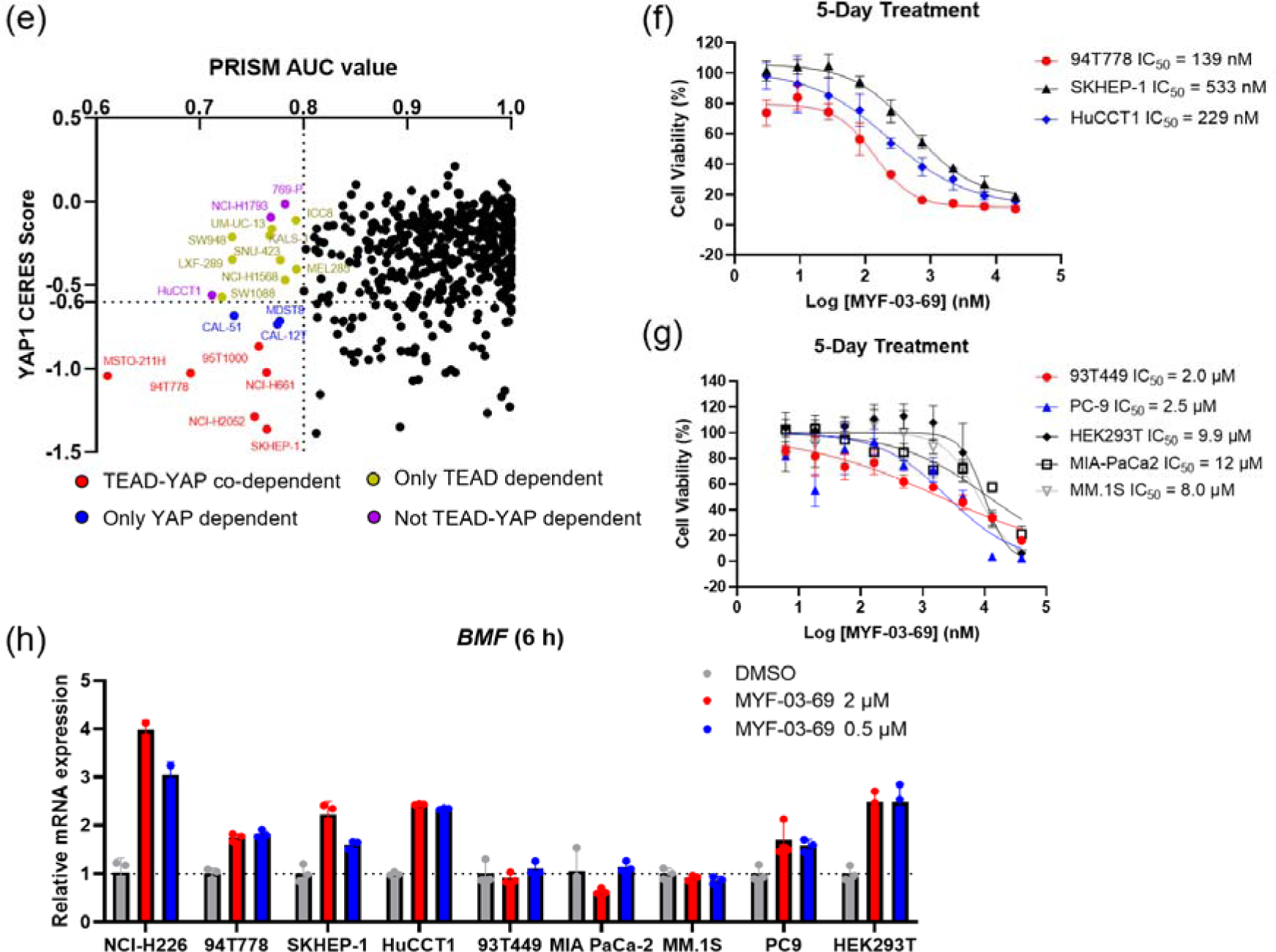
MYF-03-69 selectively inhibits proliferation of cancer cells with YAP or TEAD dependency regardless of lineage. (**a**) Antiproliferation IC_50_ curve of MYF-03-69 on three mesothelioma cell lines (NCI-H226, MSTO-211H and NCI-H2452) and a normal noncancerous mesothelium cell line (Met-5A). (**b**) Antiproliferation IC_50_ curve of MYF-03-69 and MYF-03-69-NC in 3D cell culture. (**c-d**) Cell cycle arrest induced by MYF-03-69, but not MYF-03-69-NC. Cells were treated for 48 hours at indicated doses. (**e**) PRISM profiling across a broad panel of cell lineages. 903 cancer cells were treated with MYF-03-69 for 5 days. The viability values were measured at 8-point dose manner (3-fold dilution from 10 μM) and fitted a dose-response curve for each cell line. Area under the curve (AUC) was calculated as a measurement of compound effect on cell viability. CERES score of YAP1 or TEADs from CRISPR (Avana) Public 21Q1 dataset (DepMap) were used to estimate gene-dependency. Cell lines without CERES Score of YAP1 were excluded from the figure. The CERES Score of most dependent TEAD isoform was used to represent TEAD dependency. Cell lines with a dependency score less than -0.6 were defined as the dependent cell lines. (**f**) Antiproliferation curves of cell lines that are sensitive to MYF-03-69 treatment besides mesothelioma. (**g**) Antiproliferation curves of cell lines that are insensitive to MYF-03-69 treatment. (**h**) *BMF* expression level after treatment with 0.5 or 2 μM MYF-03-69 over 6 hours in different cells.

### MYF-03-69 inhibits YAP or TEAD-dependent cancer cells beyond mesothelioma

To further investigate whether inhibition by MYF-03-69 was selectively lethal to YAP/TEAD-dependent cancers, 903 barcoded cancer cell lines were screened using the PRISM assay^23^. As shown in **Figure 5e**, a small portion of cell lineages exhibited vulnerability (**Supplementary Dataset 3**). Correlation analysis reveals that the dependency scores of *TEAD1* and *YAP1* according to genomic knockout dataset (DepMap) provided the highest correlation with the sensitivity profile (**Figure S9c, Supplementary Dataset 4**). This is followed by *TP53BP2*, a gene that is also involved in Hippo pathway as activator of TAZ^32^. For example, when we used a threshold of AUC ≤ 0.8 for the sensitivity to MYF-03-69, 33 cell lines were selected and the majority of which are either YAP or TEADs dependent cells, as suggested by CERES scores (**Figure 5e**). These include YAP/TEAD co-dependent cells (red dots), YAP dependent cells (blue dots), and TEAD dependent cells (yellow dots). Next, to verify the antiproliferative activity, we chose three sensitive cell lines (94T778, SKHEP-1 and HuCCT1) and three insensitive cell lines (93T449, MIA PaCa-2 and MM.1S) indicated by PRISM screening, as well as two additional cell lines (PC9, HEK293T) that were not included in PRISM screening panel to test in 5-day antiproliferation assay. As shown in **Figure 5f**, liposarcoma cell 94T778, hepatic adenocarcinoma cell SKHEP-1 and cholangiocarcinoma cell HuCCT1 were inhibited with nanomolar IC_50_. In contrast, liposarcoma cell 93T449, pancreatic ductal adenocarcinoma cell MIA PaCa-2 and myeloma cell MM.1S were barely inhibited (**Figure 5g**). Since proapoptotic gene *BMF* was known to be released from repression upon TEAD inhibition^10^, we examined *BMF* mRNA levels after 6-hour treatment with 0.5 and 2 µM MYF-03-69 on both sensitive and insensitive cells. As expected, *BMF* levels increased in three sensitive cells, but remained unchanged in three insensitive cells (**Figure 5h**). We also noted the upregulation of *BMF* in PC9 and HEK293T cells which are resistant to TEAD inhibition (**Figure 5g**-**h**). A heterogeneity of signaling pathway dependency, especially EGFR signaling and YAP1 activation in PC9 cells might account for the insensitivity to TEAD inhibition alone. Taken together, we demonstrate that TEAD inhibition could be an exploitable vulnerability across multiple malignant tumor models besides mesothelioma and that upregulation of *BMF* gene is a common phenomenon in those cancer cells with sensitive antiproliferative response to TEAD inhibitor.

### MYF-03-176 exhibits significant antitumor efficacy in NCI-H226 xenograft mouse model through oral administration

In order to demonstrate therapeutic potential of covalent TEAD palmitoylation disruptor, extensive medicinal chemistry efforts were undertaken on this Y-shaped scaffold, leading to a more potent and more importantly, orally bioavailable MYF-03-69 analog MYF-03-176 (**Figure 6a**). Before we administrated the mice with MYF-03-176, we conducted a head-to-head comparison on the activity with MYF-03-69 as well as other reported TEAD inhibitors including K-975, compound 2 and VT103. We used a stably transfected TEAD luciferase reporter system in NCI-H226 cells to profile transcriptional effect of TEAD inhibitors. Consistently with the results of mCherry reporter system, MYF-03-69 inhibited the transcription of reporter gene at IC_50_ of 45 nM (**Figure S10a**) and interestingly MYF-03-176 was 3-fold more potent with IC_50_ of 11 nM (**Figure 6b**). Further, similar to MYF-03-69, MYF-03-176 led to a significant downregulation of YAP target genes *CTGF*, *CYR61*, *ANKRD1* and an upregulation of *BMF* (**Figure S10b**). As comparison, MYF-03-176 exhibited comparable and even better activity than K-975 and compound 2, while was less potent than VT103 as shown in **Figure 6b∼c**. The antiproliferative effect of MYF-03-176 in NCI-H226 cells was similar to VT-103 and K-975 but even stronger than MYF-03-69 and compound 2. With the support of all these data from MYF-03-176 and decent pharmacokinetics properties including low clearance and high oral bioavailability (**Figure S10c-d**), we then administrated MYF-03-176 to NCI-H226 cell line derived xenograft (CDX) mouse model. The tumor-bearing mice were randomized and orally administrated MYF-03-176 twice daily for 28 days. Significant antitumor activity with tumor regressions was observed at both 30 mg/kg (average tumor regression of 54%) and 75 mg/kg (average tumor regression of 68%). The anti-tumor activity at the two doses was comparable (*P* = 0.23, 2-way ANOVA). The 30 mg/kg dose was well tolerated with an average body weight gain comparable to vehicle control (data not shown), while at the 75 mg/kg dose, an average wight loss of 5% was observed. However, at this 75 mg/kg dose, 3 of 8 animals demonstrated 12-14% body weight loss. The weight loss recovered once drug administration was stopped (**Figure 6f**). Taking all these together, MYF-03-176 potently inhibited Hippo signaling defective MPM cells and exhibited strong antitumor effect in the human mesothelioma NCI-H226 CDX model *in vivo*, representing a promising leading compound for drug discovery.

**Figure 6.**
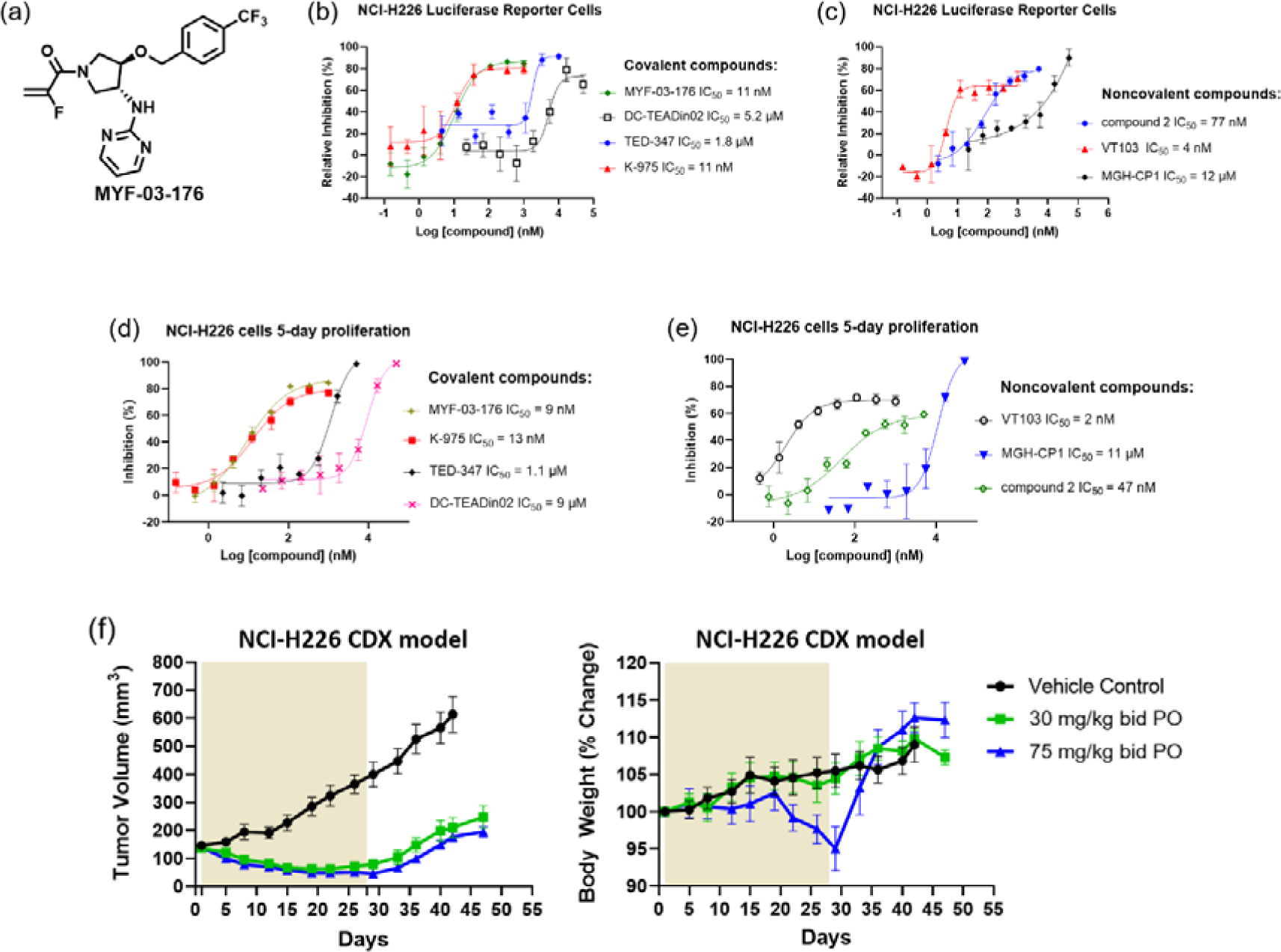
MYF-03-176 is a potent and orally bioavailable YAP-TEAD transcription inhibitor and suppresses tumor growth in mesothelioma xenograft mouse model. (**a**) Chemical structure of MYF-03-176. (**b**-**c**) Inhibitory effect of MYF-03-176 and other TEAD PBP binders on YAP-TEAD transcription in NCI-H226 luciferase reporter cells. Cells were treated for 72 hours. Data were presented as mean ± SD of n = 3 biological independent samples. (**d**-**e**) Antiproliferation effect of MYF-03-176 and other TEAD PBP binders in NCI-H226 cells. Cells were treated for 5 days. Data were presented as mean ± SD of n = 3 biological independent samples. (**f**) *In vivo* efficacy of MYF-03-176 in NCI-H226 CDX mouse model (n = 8∼9 per group).

## Discussion

As widely recognized oncogenic proteins, YAP/TAZ have emerged as potentially attractive targets for anti-cancer drug development. However, the unstructured nature of YAP/TAZ proteins renders them difficult to target by conventional occupancy-based small molecules. Therefore, the majority of compounds that are known to inhibit YAP activity are targeted at upstream stimulators of YAP/TAZ. In this context, TEADs, key components of Hippo signaling pathway that depend on YAP/TAZ binding for activation of transcriptional activity, have attracted attention. Here, the presence of a well-defined palmitate binding pocket (PBP) on TEADs suggested opportunity for small molecule inhibitor development. However, recent studies have not fully resolved the question of whether occupying TEAD PBP disrupts or stabilizes YAP-TEAD interaction. Chan et al. suggested that palmitoylation of TEAD stabilizes YAP-TEAD interaction^13b^. Holden et al. reported the reversible PBP binder compound 2 had minimal to no disruptive effect on YAP-TEAD interaction biochemically but transformed TEAD into a dominant-negative transcriptional repressor^16^. VT series compounds reported by Tang et al. dissociated endogenous YAP-TEAD according to co-IP experiment^17^, which is consistent with our result of MYF-03-69. Other data from MGH-CP1^15^ and covalent binder K-975^20^ pointed to the same conclusion while the experiments were performed with exogenous YAP or TEAD. K-975 and TED-347 were reported to disrupt protein-protein interaction between TEAD and YAP peptide in biochemical assay. However, their co-crystal structures indicated no conformation change compared with palmitoylated TEAD. Similarly, superimposition of co-crystal structures of TEADs with MGH-CP1, compound 2, VT105, MYF-03-69 and their corresponding palmitoylated TEADs did not reveal any obvious conformation change or side chain move that might affect YAP binding.

Taken all these together, the transcriptional inhibitory effect of these TEAD PBP binders might result from disruption of certain dynamic process of TEAD lifecycle. Recently, several reports elaborated that YAP/TAZ and TEAD undergo lipid-lipid phase separation (LLPS) during transcription process^33^. One hypothesis is the alkyl chain of palmitoyl might escape from TEAD PBP and expose to outside when forming such functional transcription compartment. The hydrophobic nature of the lipid chain may play a role in organizing the disordered and hydrophobic region of YAP. Thus, replacement of palmitate by these PBP binders leads to incapability of TEAD to organize into transcription machinery, though the direct binding to YAP peptide may not being affected *in vitro*. The underlying mechanism is remained to be uncovered in the future.

MYF-03-69 is a covalent compound that we designed to target the conserved cysteine on TEAD, the site of palmitoylation. The starting point for MYF-03-69 was a fragment hit that we identified through screening of a biased covalent fragment library and further optimized using structure-based strategy. Unlike screening reversible fragments which often require significant follow-up efforts to determine binding site, the MS-guided covalent tethering method gives unambiguous information of labeling site in a cost and time efficient way. Unlike previously reported compounds that engage only the hydrophobic PBP, our optimized inhibitor was developed to exploit interactions with not only the PBP but a hydrophilic binding pocket that we have identified during this study. Thus, our optimized inhibitor MYF-03-69 is a Y-shaped molecule which efficiently occupies two pockets, as well as covalently binds the conserved cysteine, which contributed significantly to both potency and specificity. Therefore, we speculated that MYF-03-69 may be employed as a useful tool to interrogate the process of how PBP binder affect TEAD homeostasis. We also propose that further optimization of the basic Y-shaped molecule reported here may yield paralog-selective TEAD inhibitors.

Aberrant regulation of Hippo-YAP/TAZ-TEAD signaling axis has been recognized as driver for genesis and development of multiple cancers, especially in mesothelioma. Besides the Hippo pathway, diverse signaling pathways such as Wnt, TGF-β, EGFR and Hedgehog pathways can also potentiate the activation of YAP/TAZ. Given such essentiality, it is important to ascertain what portion of these YAP/TAZ activated cancers are vulnerable to TEAD inhibition. The pan-TEAD selective nature of MYF-03-69 allowed us to examine this question at the TEAD family level using a panel of more than 900 cancel cell lines. Our results demonstrated that MYF-03-69 exhibited selective antiproliferative effect on YAP or TEAD dependent cancer cells from different lineages including mesothelioma, liver cancer, liposarcoma, and lung cancer. Strong antitumor efficacy in human mesothelioma NCI-H226 CDX model has been achieved with an orally bioavailable compound MYF-03-176 in the same scaffold, which is ready to be tested in various cancer models. Moreover, we noted that increased expression of a proapoptotic *BMF* gene correlates with response to TEAD inhibition in some cancer cell lines.

Collectively, this study provides evidence that MYF-03-69 represents a potent, covalent, cysteine-targeted pan-TEAD inhibitor that disrupts YAP-TEAD association and affects transcription in covalent binding-dependent manner. Given its “mild” covalent warhead and low reactivity across proteome, we nominate MYF-03-69 as a viable lead for drug discovery for not only mesothelioma, but other YAP or TEAD dependent cancers such as liver cancer, liposarcoma, and lung cancer.

## Acknowledgments

The authors would like to thank the following for valuable help with this study: Dr. Milka. Kostic for her advice and editing for this manuscript; Jim Sun at the NMR facility of the Dana-Farber Cancer Institute for his assistance on ^1^H NMR data collection; Zachary Herbert from the Molecular Biology Core Facility at the Dana-Farber Cancer Institute for the sequencing services; Kara Soroko and Jessica Sarro from Dana-Farber Cancer Institute for the animal study services. J.D.M. acknowledges this work was supported by the Hale Family Center for Pancreatic Cancer Research.

## Author contributions

N.S.G. and T.Z. conceived the project. M.F. performed the compound synthesis and structure determination with help from Y.L., W.J. and Z.H.. W.L. executed biological experimental research with help from Y.G., N.K., J.J. and J.T.. J.C. executed computational modeling and RNA sequencing analysis. S.B.F. and J.A.M. executed and analyzed protein mass spectrum. H.S., E.A.G., J.L., K.S. and S.D.P. executed protein expression, purification, crystallization and structural determination. P.C.G. managed the animal study and analyzed the results. M.K. and J.D.M. performed the SLC-ABPP profiling and analysis. A.S.B., M.G.R., M.M.R. and J.A.R. executed PRISM screening and analysis. M.F., W.L., T.Z., J.C. and N.S.G. co-wrote the paper. All authors edited the manuscript.

## EXPERIMENTAL DETAILS, METHODS AND CHARACTERIZATION DATA

### 1.1 Cloning

The stretch of residues 209-424 of human TEAD1, residues 220-450 of human TEAD2, residues 119-436 of human TEAD3, residues 216-434 of human TEAD4, were inserted into the pET28PP (N-terminal His 3C tag) vector.

### 1.2 Protein expression and purification

The N-terminal His tag construct of human TEAD1 (residues 209-424) was overexpressed in E. coli BL21 (DE3) and purified using affinity chromatography and size-exclusion chromatography. Briefly, cells were grown at 37°C in TB medium in the presence of 50 μg/ml of kanamycin to an OD of 0.8, cooled to 17°C, induced with 500 μM isopropyl-1-thio-D-galactopyranoside (IPTG), incubated overnight at 17°C, collected by centrifugation, and stored at -80°C. Cell pellets were lysed in buffer A (25 mM HEPES, pH 7.5, 200 mM NaCl, 5% glycerol, 7 mM mercapto-ethanol, and 20 mM Imidazole) using Microfluidizer (Microfluidics), and the resulting lysate was centrifuged at 30,000g for 40 min. Ni-NTA beads (Qiagen) were mixed with cleared lysate for 30 min and washed with buffer A. Beads were transferred to an FPLC-compatible column, and the bound protein was washed further with buffer A for 10 column volumes and eluted with buffer B (25 mM HEPES, pH 7.5, 200 mM NaCl, 5% glycerol, 7 mM mercapto-ethanol, and 400 mM Imidazole). The eluted sample was concentrated and purified further using a Superdex 200 16/600 column (Cytiva) in buffer C containing 20 mM HEPES, pH 7.5, 200 mM NaCl, 5% glycerol, 0.5 mM TCEP and 2 mM DTT. HRV 3C protease was added to TEAD1 containing fractions and incubated overnight at 4°C, followed by passing through Ni-NTA column to remove His-tag and 3C protease. The flow-through fractions of the second Ni-NTA column, containing cleaved TEAD1, was concentrated to ∼9mg/mL and stored in -80°C. The N-terminal His tag construct of human TEAD2 (residues 220-450), TEAD3 (residues 119-436) and TEAD4 (residues 216-434) were purified as TEAD1.

#### 2 Mass spectrometry analysis

TEAD2 protein (5 µg) was treated with DMSO or a 10-fold molar excess of MYF-03-69 and analyzed by LC-MS using an HPLC system (Shimadzu, Marlborough, MA) interfaced to an LTQ ion trap mass spectrometer (ThermoFisher Scientific, San Jose, CA). Proteins were desalted for 4 minutes on column with 100% A, and eluted with an HPLC gradient (0-100% B in 1 minute; A=0.2M acetic acid in water; B=0.2M acetic acid in acetonitrile). The mass spectrometer was programmed to acquire full scan mass spectra (*m/z* 300-2000) in profile mode (spray voltage = 4.5 kV). Mass spectra were deconvoluted using MagTran software version 1.03b2^1^. To analyze the site of modification, DMSO or MYF-03-69 treated proteins were first captured on SP3 beads^2^ by adding an equal volume of acetonitrile and washed (2x with 70% acetonitrile, 1x with 100% acetonitrile). Beads were then resuspended with 100 mM ammonium bicarbonate containing 0.1% Rapigest (Waters, Milford, MA). Proteins were reduced with 10 mM DTT for 30 minutes at 56°C, alkylated with 22.5 mM iodoacetamide for 30 minutes at room temperature, and then digested overnight with trypsin at 37 °C. Rapigest was cleaved according to the manufacturer’s instructions, and peptides were desalted by C18 and dried by vacuum centrifugation. Peptides were reconstituted in 50% acetonitrile, 1% formic acid, 100 mM ammonium acetate, and analyzed by CE-MS using a ZipChip CE-MS instrument and autosampler (908 devices, Boston, MA) interfaced to a QExactive HF mass spectrometer (ThermoFisher Scientific). Peptides were resolved at 500V/cm using an HR chip with a background electrolyte consisting of 50% acetonitrile with 1% formic acid. The mass spectrometer was operated in data dependent mode, and subjected the 5 most abundant ions in each MS scan (*m/z* 300-2000, 60K resolution, 3E6 target, 100 ms max fill time) to MS/MS (15K resolution, 1E5 target, 100 ms max fill time). Dynamic exclusion was enabled with a repeat count of 1 and an exclusion duration of 5 seconds. Raw mass spectrometry data files were converted to .mgf using multiplierz software^3^ and searched against a forward-reverse human refseq database using Mascot version 2.6.2. Search parameters specified fixed carbamidomethylation of cysteine, variable methionine oxidation, and variable MYF-03-69 modification of cysteine. MYF-03-69 modified spectra were examined and figures prepared using mzStudio software^4^. Inhibitor related ions in MS/MS spectra were identified as described^5^.

### 3 Docking

Docking of MYF-01-37 was performed with covalent docking protocol from Schrodinger suite software (release 2019-02) with default parameters in TEAD2 structure (PDB code: 5HGU) exporting 5 poses per molecule. Both stereoisomers were docked. Top scoring pose was chosen to illustrate the binding pose. The pdb structure was processed and optimized with protein preparation protocol with default setting in Schrodinger suite.

### 4.1 Crystallization

Using Formulatrix NT8 and RockImager and ArtRobbins Phoenix liquid handlers, a 100 nL sample of 300 µM TEAD1 that was preincubated for 1 hour with 600 µM MYF-03-69 was dispensed in an equal volume of crystallization buffer (3M NH4SO4 and 0.1 M Tris pH 9.0) and incubated against 25 µL of crystallization buffer in a 384-well hanging-drop vapor diffusion microtiter plate at 20 °C for three days.

### 4.2 Data collection and structure determination

Diffraction data were collected at beamline 24ID-E of the NE-CAT at the Advanced Photon Source (Argonne National Laboratory). Data sets were integrated and scaled using XDS^6^. Structures were solved by molecular replacement using the program Phaser^7^ and available search models from the PDB. Iterative manual model building and refinement using Phenix^8^ and Coot^9^ led to a model with excellent statistics.

#### 5 Gel-based palmitoylation studies

1 μM TEADs-YBD recombinant protein was incubated with inhibitors at the indicated concentrations at 37 °C for 2 h followed by the addition of palmitoyl alkyne-coenzyme A (Cayman chemical, no. 15968) in a total volume of 50 μL. After 30 min reaction, 5 μL 10%SDS were added and 5 μL click reagents were added to start click reaction as previously reported^10^. After another 1 h, 4x loading buffer were added to the reaction mixture and the samples subjected for western blot analysis. IRDye 800CW Streptavidin (LI-COR, no. 92632230) and His-Tag Mouse mAb (Cell Signaling, no. 2366S) was used for biotin detection and His-tag detection. The blots were imaged on Odyssey CLx Imager (LI-COR).

### 6.1 Sample preparation for SLC-ABPP

Samples for whole cysteine profiling were prepared as previously described^11^. Briefly, frozen cell pellets from H226 cells were lysed using PBS (pH 7.4). Samples were further homogenized, and DNA was sheared using sonication with a probe sonicator (20 x 0.5 s pulses) at 4°C. Total protein was determined using a BCA assay and cell lysates were used immediately for each experiment. Depending on the experiment, 50 μg of total cell extract was aliquoted for each TMT channel for further downstream processing. Excess DBIA, along with disulfide bonds were quenched and reduced using 5 mM dithiothreitol (DTT) for 30 min in the dark at room temperature. Subsequently reduced cysteine residues were alkylated using 20 mM iodoacetamide for 45 min in the dark at room temperature. To facilitate removal of quenched DBIA and incompatible reagents, proteins were precipitated using chloroform/methanol. Briefly, to 100 μL of each sample, 400 μL of methanol were added, followed by 100 μL of chloroform with thorough vortexing. Next, 300 μL of HPLC grade water were added, and the samples were mixed to facilitate precipitation. Samples were centrifuged at maximum speed (14,000 rpm) for 3 min at room temperature, the aqueous top layer was removed, and the samples were washed additionally three times with 500 μL of methanol. Protein pellets were re-solubilized in 200 mM 4-(2-hydroxyethyl)-1-piperazinepropanesulfonic acid (EPPS) at pH 8.5 and digested using LysC and trypsin (1:100, enzyme to protein ratios) overnight at 37°C using a ThermoMixer set to 1200 rpm. The next day samples were labeled with TMT reagents or stored at -80°C until further use.

### 6.2 TMT Labeling for SLC-ABPP

Digested peptides containing DBIA conjugated-cysteines were labeled using TMTPro 16-plex reagents as previously described^12^. Briefly, peptides were labeled at a 1:2 ratio by mass (peptides to TMT reagents) for 1 hr with shaking at 1,200 rpm. To equalize protein loading, ∼ 2 μg of each sample were aliquoted, and a 60-min quality control analysis (ratio check) was performed using SPS-MS3. Excess TMT reagent was quenched with hydroxylamine (0.3% final concentration) for 15 min at room temperature. Next, samples were mixed 1:1 across all TMT channels and the pooled sample was dried using a Speedvac to ensure all acetonitrile was removed.

### 6.3 Cysteine peptide enrichment using streptavidin in SLC-ABPP

Pierce streptavidin magnetic beads were washed with PBS pH 7.4 prior to use. To each TMT labeled pooled sample, 100 μL of a 50% slurry of streptavidin beads were added. Samples were further topped up with 1 mL of PBS in a 2 mL Eppendorf tube. Samples and beads were incubated overnight at 4 °C to enrich for TMT-labeled DBIA conjugated cysteine peptides. Following enrichment, the beads were placed onto a magnetic rack and allowed to equilibrate for 5 min. Beads were washed to remove non-specific binding using the following procedure: 3 x 1 mL of PBS pH 7.4, 1 x 1 mL of PBS with 0.1% SDS pH 7.4 and finally 3 x 1 mL HPLC grade water. Beads were resuspended using a pipette between washes and placed on the magnet between each wash. To elute cysteine containing peptides, 500 μL of 50% acetonitrile with 0.1% trifluoracetic acid (TFA) were added, and the beads were mixed at 1,000 rpm for 15 min at room temperature. Eluted peptides were transferred to a new tube, and the beads were additionally washed with 200 μL of 50% acetonitrile with 0.1% TFA and combined. Cysteine containing peptides were dried to completion using a Speedvac and were stored at -80°C.

### 6.4 Desalting cysteine-containing peptides in SLC-ABPP

TMT-labeled cysteine-containing peptides were resuspended using 200 μL of 1% formic acid (FA) and were desalted using StageTips as previously described^12a^. Briefly, eight 18-guage cores were packed into a 200 μL pipette tip and were passivated and equilibrated using the following solutions: 100 μL of 100% methanol, 70% acetonitrile 1% FA, and 5% acetonitrile 5% FA. Peptides were loaded and were washed with 0.1% FA and eluted using 150 μL of 70% acetonitrile 1% FA and dried to completion using a Speedvac. Enriched peptides samples were next resuspended in 5-10 μL of 5% acetonitrile 5% FA and between 50 and 100% of the sample was injected for analysis using LC-RTS-SPS-MS3.

### 6.5 Mass spectrometry and real-time searching in SLC-ABPP

All SLC-ABPP mass spectrometry data were acquired using an Orbitrap Fusion Eclipse mass spectrometer in-line with a Proxeon NanoLC-1200 UPLC system. Peptides were separated using an in-house 100 µm capillary column packed with 35 cm of Accucore 150 resin (2.6 μm, 150 Å) (ThermoFisher Scientific) using 210 min gradients from 4 to 24% acetonitrile in 0.125% formic acid per run. Eluted peptides were quantified using the synchronous precursor selection (SPS-MS3) method for TMT quantification. Briefly, MS1 spectra were acquired at 120K resolving power with a maximum of 50 ms ion injection in the Orbitrap. MS2 spectra were acquired by selecting the top 10 most abundant features via collisional induced dissociation (CID) in the ion trap using an automatic gain control (AGC) setting of 15K, quadrupole isolation width of 0.5 m/z and a maximum ion accumulation time of 50 ms. These spectra were passed in real time to the external computer for online database searching. Intelligent data acquisition (IDA) using real-time searching (RTS) was performed using Obiter^13^. Cysteine-containing peptides or whole proteome peptide spectral matches were analyzed using the Comet search algorithm (release_2019010) designed for spectral acquisition speed^14^. The same forward- and reversed-sequence human protein databases were used for both the RTS search and the final search (Uniprot). The RTS Comet functionality has been released and is available here: http://cometms.sourceforge.net/. Real-time access to spectral data was enabled by the Thermo Scientific Fusion API (https://github.com/thermofisherlsms/iapi). Next, peptides were filtered using simple initial filters that included the following: not a match to a reversed sequence, maximum PPM error of <50, minimum XCorr of 0.5, minimum deltaCorr of 0.10 and minimum peptide length of 7. If peptide spectra matched to above criteria, an SPS-MS3 scan was performed using up to 10 *b-* and *y-type* fragment ions as precursors with an AGC of 200K for a maximum of 200 ms with a normalized collision energy setting of 55 (TMTPro 16-plex)^12b^.

### 6.6 Mass spectrometry data analysis in SLC-ABPP

All acquired data were searched using the open-source Comet algorithm (release_2019010) using a previously described informatics pipeline^15^. Spectral searches were done using a custom FASTA-formatted database which included common contaminants, reversed sequences (Uniprot Human, 2014) and the following parameters: 50 PPM precursor tolerance, fully tryptic peptides, fragment ion tolerance of 0.9 Da and a static modification by TMTPro (+304.2071 Da) on lysine and peptide N termini. Carbamidomethylation of cysteine residues (+57.021 Da) was set as a static modification while oxidation of methionine residues (+15.995 Da) and DBIA on cysteine residues (+239.262) was set as a variable modification. Peptide spectral matches were filtered to a peptide false discovery rate (FDR) of less than 1% using linear discriminant analysis employing a target-decoy strategy. Resulting peptides were further filtered to obtain a 1% protein FDR at the entire dataset level (including all plexes per cell line), and proteins were collapsed into groups. Cysteine-modified peptides were further filtered for site localization using the AScore algorithm with a cutoff of 13 (p<0.05) as previously described^15b^. Overlapping peptide sequences that were generated from different charge states, elution times and tryptic termini were grouped together into a single entry. A single quantitative value was reported, and only unique peptides were reported. Reporter ion intensities were adjusted to correct for impurities during synthesis of different TMT reagents according to the manufacturer’s specifications. For quantification of each MS3 spectrum, a total sum signal-to-noise of all reporter ions of 160 (TMTPro) was required. Lastly, peptide quantitative values were normalized so that the sum of the signal for all proteins in each channel was equal to account for sample loading differences (column normalization).

### 6.7 SLC-ABPP competition ratio calculation

Cysteine site-specific engagement was assessed by the blockage of DBIA probe labeling. Peptides that showed >95% reduction in TMT intensities in the electrophile-treated samples were assigned a maximum ratio of 20 for graphing purposes with preserved ranking. TMT reporter ion sum-signal-to-noise for each SLC-ABPP experiment was used to calculate the competition ratios by dividing the control channel (DMSO) by the electrophile treated channel. Replicate measurements were averaged and reported as a single entry. To avoid false positives, sites with large coefficients of variation had the highest replicate CR values removed before averaging as previously described^11, 16^.

### 7.1 Cell culture

HEK293T (ATCC, no. CRL-3216), NCI-H226 (ATCC, no. CRL-5826), MSTO-211H (ATCC, no. CRL-2081), NCI-H2452 (ATCC, no. CRL-5946) mesothelioma cells, MeT-5A (ATCC, no. CRL-9444) mesothelium cells, 94T778 cells (ATCC, no. CRL-3044), 93T449 cells (ATCC, no. 3043), MIA PaCa-2 cells (ATCC, no. CRM-CRL-1420), MM.1S (ATCC, no. CRL-2974) and SKHEP-1 cells (ATCC, no. HTB-52) were obtained from American Type Culture Collection and cultured as recommended. HuCCT1 cells were gifts from Bardeesy lab in Massachusetts General Hospital. PC9 cells were gifts from Pasi lab in Dana-Farber Cancer Institute. Cells were negative for mycoplasma using MycoAlert mycoplasma detection kit (LONZA, no. LT07-418).

### 7.2 Cell proliferation assay

For 2D adherent cell viability experiment, the cells were seeded at 384-well plate (Corning, no. 3570) at the density of 200 cells/well. The next day, compounds were added using Janus workstation (PerkinElmer). After 5 days treatment, the cell viability was measured by CellTiter-Glo kit (Promega, no. G7570) as the manufacturer recommended. For 3D spheroid assays, NCI-H226 and MSTO-211H cells were plated at the density of 200 cells/well in Ultra-low Attachment (ULA) plate (S-bio, no. MS-9384WZ) without or with 5% Matrigel matrix (Corning, no. 356231) respectively. The cell viability was measured using 3D CellTiter-Glo kit (Promega, no. G9681). The luminescent signal was collected on EnVision plate reader (PerkinElmer). The GR_50_ values were calculated as previously described^17^.

### 7.3 Cell cycle analysis

Cells were plated in 6-well plate (Corning, no. 3506) and treated by MYF-03-69 at indicated concentrations. Then cells were harvested and fixed in cold 70% ethanol overnight. Next day, the samples were treated with 100 μg/mL RNAase A (Life Technologies, no. EN0531) and stained by 50 μg/mL propidium iodide solution (Life Technologies, no. P3566). After incubation at room temperature for 30 min, the samples were subjected to assessment using Guava flow cytometer. The data were further analyzed in Flowjo software.

#### 8 PRISM screening and data analysis

Up to 931 barcoded cell lines in pools of 20-25 were thawed and plated into 384-well plates (1250 cells/well for adherent cell pools, 2000 cells/well for suspension or mixed suspension/adherent cell pools) containing compound (top concentration: 10 µM, 8-point, threefold dilution). All conditions were tested in triplicate. Cells were lysed after 5 days of treatment and mRNA based Luminex detection of barcode abundance from lysates was carried out as described previously^18^. Luminex median fluorescence intensity (MFI) data was input to a standardized R pipeline (https://github.com/broadinstitute/prism_data_processing) to generate viability estimates relative to vehicle treatment for each cell line and treatment condition, and to fit dose-response curves from viability data. Correlation analysis was also performed in the R pipeline mentioned above.

#### 9 TEAD reporter assays

The TEAD transcriptional vector (TBS-mCherry) was used to produce lentivirus as previously reported in HEK293T cells^19^. The virus was collected at 48 h and 72 h post transfection and concentrated using Lenti-X concentrator (Takara Bio, no. 631231). Then NCI-H226 cells were transduced using the concentrated virus in the presence of 8 μg/mL polybrene (MilliporeSigma, no. TR1003G). The positively transduced cells were further sorted by GFP. For reporter assays, the selected cells were plated in black 384-well plate (Corning, no. 4514) at the density of 1,000 cells/well. The next day, compounds were added at indicated concentrations using Janus workstation. After 72 h incubation, the cells were stained by Hoechst 33342 (Life Technologies, no. 62249). The mCherry and Hoechst signals were read using Acumen high content imager (TTP Labtech).

For the luciferase reporter assay, NCI-H226 cells were transduced with 250 uL TEAD luciferase reporter lentivirus (BPS Biosciences, no. 79833) in the presence of 8 μg/mL polybrene (MilliporeSigma, no. TR1003G). Cells were then selected by 2 μg/mL puromycin. The selected cells were seeded at the density of 1,000 cells/well and treated with compounds at indicated concentrations. After three days treatment, ONE-Glo™ luciferase assay kit (Promega, no. E6110) and CellTiter-Glo kit (Promega, no. G9681) were used according to manufacturer instructions. The TEAD transcriptional activity was then calculated by normalizing luciferase signal normalized with relative cell viability.

#### 10 Lipid displacement assay

The HEK-293T cells were plated in six-well plate (Corning, no. 3506) and transfected with Myc-TEAD4 vector the next day. The Myc-TEAD4 was a gift from Kunliang Guan^20^ (Addgene plasmid # 24638; http://n2t.net/addgene:24638; RRID: Addgene_24638). Then transfected cells were treated with 50 μM alkyne palmitic acid (Click Chemistry Tools, cat. no. 1165-5) in the presence of TEAD inhibitors at indicated concentrations next day. After 24-hour incubations, cells were collected and lysed in RIPA buffer with proteasome inhibitor. The cleared supernatant was then subjected to click reactions using same procedures in the gel-based palmitoylation assay. Palmitoylation levels were detected by immunoblot using streptavidin antibody (LI-COR, no. 92632230).

#### 11 RT-PCR studies

The cells were plated in six-well plate (Corning, no. 3506) and treated with compounds the next day. The total RNA was extracted using RNeasy Plus Mini Kit (Qiagen, no. 74134). Then 500 ng purified RNA was used to synthesize cDNA by SuperScript III First-Strand Synthesis System Kit (Life Technologies, no. 18080051). The following TaqMan probes were used for follow-up RT-PCR reactions: CTGF (Hs00170014_m1), CYR61 (Hs00155479_m1), GAPDH (Hs02786624_g1), BMF (Hs00372937_m1), IGFBP3 (Hs00181211_m1), KRT34 (Hs02569742_s1) and NPPB (Hs00173590_m1).

#### 12 Co-immunoprecipitation (Co-IP) studies

The cells were plated in 10 cm dish (Corning, no. 430293) and treated with compounds at indicated concentrations the next day. The Co-IP experiments were conducted using Dynabeads Co-Immunoprecipitation Kit (Life Technologies, no. 14321D) according to the manufacturer instructions.

### 13.1 RNA-Seq study: Sample Treatment

The NCI-H226 cells were plated in the 10 cm dish and treated with MYF-03-69 at indicated concentrations with biological triplicates for 6 hours next day. The RNA was extracted using RNeasy plus mini kit (Qiagen, cat no.74134) according to the manufacturer instructions.

### 13.2 RNA-Seq study: Library Preparation and Sequencing

Libraries were prepared using Roche Kapa mRNA HyperPrep strand specific sample preparation kits from 200 ng of purified total RNA according to the manufacturer’s protocol on a Beckman Coulter Biomek i7. The finished dsDNA libraries were quantified by Qubit fluorometer and Agilent TapeStation 4200. Uniquely dual indexed libraries were pooled in an equimolar ratio and shallowly sequenced on an Illumina MiSeq to further evaluate library quality and pool balance. The final pool was sequenced on an Illumina NovaSeq 6000 targeting 40 million 100bp read pairs per library at the Dana-Farber Cancer Institute Molecular Biology Core Facilities.

### 13.3 RNA-Seq study: Analysis

Sequenced reads were aligned to the UCSC hg19 reference genome assembly and gene counts were quantified using STAR (v2.7.3a)^21^. Differential gene expression testing was performed by DESeq2 (v1.22.1)^22^. RNAseq analysis was performed using the VIPER snakemake pipeline^23^. KEGG pathway enrichment analysis was performed through metascape webportal^24^.

### 13.4 RNA-Seq study: Datasets availablility

These RNA-seq datasets were deposited to BioSample database under accession# as below: SAMN19288936, SAMN19288937, SAMN19288938, SAMN19288939, SAMN19288940, SAMN19288941, SAMN19288942, SAMN19288943, SAMN19288944, SAMN19288945, SAMN19288946.

### 14.1 Efficacy study in NCI-H226 CDX model: Animals

Female 7-week-old NSG mice were purchased from The Jackson Laboratory (Bar Harbor, ME). Animals acclimated for at least 5 days before initiation of the study. All *in vivo* studies were conducted at Dana-Farber Cancer Institute with the approval of the Institutional Animal Care and Use Committee in an AAALAC accredited vivarium.

### 14.2 Efficacy study in NCI-H226 CDX model: *In vivo* studies

The NCI-H226 cells were grown in RPMI1640 media supplemented with 10% fetal bovine serum. Cells were harvested, and 5 × 10^6^ cells with 50% Matrigel (Fisher Scientific) were implanted subcutaneously in the right flank of the NSG mice. Tumors were allowed to establish to an average of 141.3 ± 24.9 mm^3^ in size before randomization using Studylog software (San Francisco, CA) into various treatment groups with 8-9 mice per group. MYF-03-176 was formulated as a suspension in 10% DMSO with 10% Tween 80 in water and dosed twice daily via oral gavage. Control treated mice received vehicle alone. Tumor volumes were determined from caliper measurements by using the formula, Tumor volume = (length × width^2^)/2. Tumor volumes and body weights were measured twice weekly. Mice were treated for 28 days, followed by measuring for re-growth of tumors.

#### 15 Chemical synthesis and characterization

**MYF-03-42**

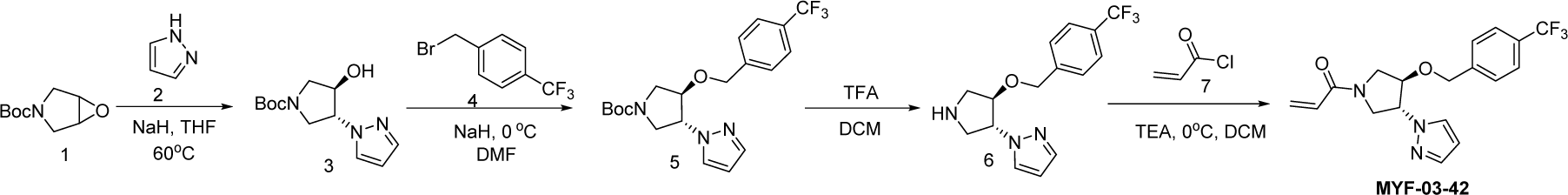

#### Step 1: Synthesis of *trans*-tert-butyl 3-hydroxy-4-(1H-pyrazol-1-yl)pyrrolidine-1-carboxylate (Compound 3)

To a suspension of 60% NaH (323mg, 8.1 mmol) in THF (20 mL) was added the solution of 1H-pyrazole (551 mg, 8.1 mmol) in THF (5 mL), the mixture was stirred at 0 °C under N_2_ for 30 minutes, and then tert-butyl 6-oxa-3-azabicyclo[3.1.0] hexane-3-carboxylate (1.0 g, 5.4 mmol) was added. The resulting mixture was heated at 60°C under N_2_ for 16 hours. After cooled down to room temperature the mixture was diluted with ethyl acetate (50 mL) and washed with water (50 mL). The organic layer was dried over anhydrous Na_2_SO_4_, filtered, concentrated and purified by flash column chromatography on silica gel (ethyl acetate in petroleum ether = 40% v/v) to afford compound **3** as solid (600 mg, yield 44%). LC-MS (ESI) m/z: 254 [M+H]^+^.

#### Step 2: Synthesis of *trans*-tert-butyl 3-(1H-pyrazol-1-yl)-4-(4-(trifluoromethyl)benzyloxy)pyrrolidine-1-carboxylate (Compound 5)

A mixture of compound **3** (300 mg, 1.18 mmol), 1-(bromomethyl)-4-(trifluoromethyl)benzene (283 mg, 1.18 mmol) and 60% NaH (47 mg, 1.18 mmol) in DMF (10 mL) was stirred at room temperature under N_2_ for 2 hours. The mixture was diluted with water (50 mL) and extracted with ethyl acetate (50 mL x 2), the combined organic was washed with brine (50 mL), dried over anhydrous Na_2_SO_4_, filtered and concentrated to obtain compound **5** as yellow oil (400 mg, yield 82%). LC-MS (ESI) m/z: 356 [M-56+H]^+^.

#### Step 3: Synthesis of 1-(*trans*-4-(4-(trifluoromethyl)benzyloxy)pyrrolidin-3-yl)-1H-pyrazole (Compound 6)

A mixture of compound **5** (300 mg, 0.72 mmol) and TFA (3 mL) in DCM (10 mL) was stirred at room temperature for 2 hours. The mixture was concentrated to leave crude compound **6** (250 mg) as yellow oil, which was used directly in the next step. LC-MS (ESI) m/z: 312[M+H]^+^.

#### Step 4: Synthesis of 1-(*trans*-3-(1H-pyrazol-1-yl)-4-(4-(trifluoromethyl)benzyloxy)pyrrolidin-1-yl)prop-2-en-1-one (MYF-03-42)

A mixture of compound **6** (70 mg, 0.22 mmol), acryloyl chloride (20 mg, 0.22 mmol), and TEA (44 mg, 0.44mmol) in DCM (10 mL) was stirred at room temperature under N_2_ for 2 hours. The mixture was concentrated, the residue was purified by prep-HPLC to afford **MYF-03-42** as yellow oil (50 mg, yield 62%). LC-MS (ESI) m/z: 366[M+H]^+^. ^1^H-NMR (400 MHz, CD_3_OD): δ (ppm) 7.74 (dd, *J* = 5.2, 2.4 Hz, 1H), 7.63 (d, *J* = 8.0 Hz, 2H), 7.56 (d, *J* = 1.6 Hz, 1H), 7.47 (d, *J* = 8.1 Hz, 2H), 6.66-6.59 (dd, *J* = 16.8, 10.4 Hz, 1H), 6.36-6.30 (m, 2H), 5.85 – 5.74 (m, 1H), 5.18 – 5.02 (m, 1H), 4.68 (d, *J* = 5.8 Hz, 2H), 4.51-4.42 (m, 1H), 4.28-4.08 (m, 2H), 4.01-3.93 (m, 1H), 3.83-3.65 (m, 1H).

**MYF-02-111**

**MYF-02-111** was synthesized through the same route as **MYF-03-42** except 1-(bromomethyl)-3-(trifluoromethyl)benzene was used in **Step 2** instead of 1-(bromomethyl)-4-(trifluoromethyl)benzene. **MYF-02-111** was obtained as light yellow oil. LC-MS (ESI) m/z: 366[M+H]^+^. ^1^H NMR (500 MHz, DMSO-*d_6_*) δ 7.87 (ddd, J = 9.3, 2.3, 0.7 Hz, 1H), 7.69 – 7.54 (m, 4H), 7.53 – 7.48 (m, 1H), 6.61 (dd, J = 16.8, 10.3 Hz, 1H), 6.30 (dt, J = 3.4, 2.1 Hz, 1H), 6.17 (dt, J = 16.7, 2.5 Hz, 1H), 5.70 (td, J = 9.9, 2.3 Hz, 1H), 5.13 (ddt, J = 33.6, 7.8, 3.4 Hz, 1H), 4.66 (d, J = 4.0 Hz, 2H), 4.39 (ddd, J = 25.5, 4.1, 1.7 Hz, 1H), 4.19 – 4.09 (m, 0.5H), 4.07 – 3.87 (m, 1.5H), 3.86 – 3.70 (m, 1.5H), 3.52 (dd, J = 13.0, 4.0 Hz, 0.5H).

**MYF-03-69**

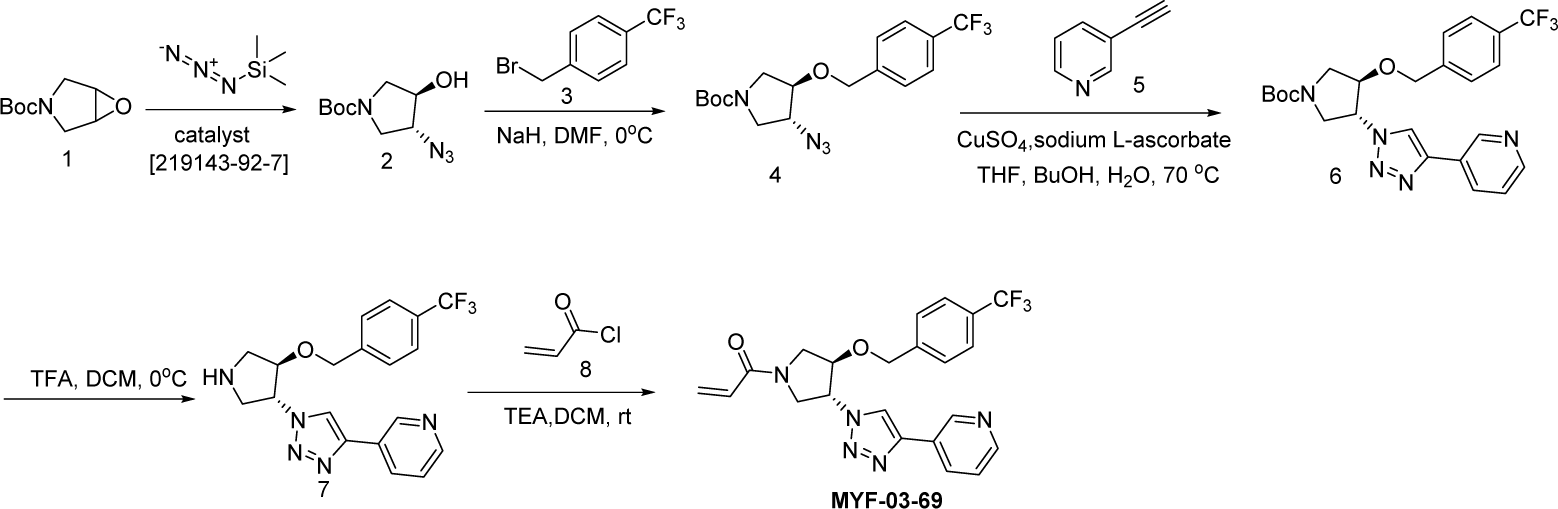

#### Step 1: Synthesis of (*3R*, *4R*)-tert-butyl 3-azido-4-hydroxypyrrolidine-1-carboxylate (Compound 2)

A mixture of tert-butyl 6-oxa-3-azabicyclo[3.1.0]hexane-3-carboxylate (4 g, 21.6 mmol), TMSN_3_ (2.664 g, 23.2 mmol) and chiral catalyst (*1S*,*2S*)-(-)-[1,2-cyclohexanediamino-N,N’-bis(3,5-di-t-butylsalicylidene)]chromium(III) chloride (328 mg, 0.42 mmol) was stirred at rt under N_2_ overnight. The reaction mixture was treated with MeOH (60 mL) and K_2_CO_3_ (1.788 g, 12.8 mmol) and continued to stir at rt for 5 hours. The reaction mixture was diluted with ethyl acetate (300 mL), washed with water (300 mL x 2), dried over anhydrous Na_2_SO_4_, concentrated and purified by flash column chromatography on silica gel (ethyl acetate in petroleum ether = 30% v/v) to obtain the compound **2** as clear oil (3.5g, 96% e.e., yield 71%). LC-MS (ESI) m/z: 129 [M+H-100]^+^.

#### Step 2: Synthesis of (*3R*, *4R*)-tert-butyl 3-azido-4-(4-(trifluoromethyl)benzyloxy)pyrrolidine-1-carboxylate (Compound 4)

A mixture of compound **2** (3 g, 13.1 mmol), 1-(bromomethyl)-4-(trifluoromethyl)benzene (3.1 g, 13.1 mmol) and 60% NaH (0.6 g, 15.7 mmol) in DMF (20 mL) was stirred at 0 °C under N_2_ for 6 hours. The reaction mixture was diluted with water (200 mL) and extracted with ethyl acetate (200 mL), the organic was washed with water (100 mL), dried over anhydrous Na_2_SO_4_, concentrated and purified by flash column chromatography on silica gel (ethyl acetate in petroleum ether = 20% v/v) to obtain compound **4** as oil (3.8g, yield 75%). LC-MS (ESI) m/z: 287[M+H-100]^+^.

#### Step 3: Synthesis of (*3R*, *4R*)-tert-butyl 3-(4-(pyridin-3-yl)-1H-1,2,3-triazol-1-yl)-4-(4-(trifluoromethyl)benzyloxy)pyrrolidine-1-carboxylate (Compound 6)

A mixture of compound **4** (3.8 g, 9.8 mmol), 3-ethynylpyridine (1.216 g, 11.8 mmol), CuSO_4_ (244 mg, 0.98 mmol) and, sodium L-ascorbate (388 mg, 1.96 mmol) in THF (20 mL), H_2_O (20 mL) and ^n^BuOH (20 mL) was stirred at 70 °C under N_2_ overnight. The reaction mixture was concentrated in vacuum, the residue was diluted with ethyl acetate (600 mL), washed with water (400 mL), dried over anhydrous Na_2_SO_4_, concentrated and purified by flash column chromatography on silica gel (ethyl acetate in petroleum ether = 90% v/v) to obtain compound **6** as yellow solid (4g, yield 83%). LC-MS (ESI) m/z: 490[M+H]^+^.

#### Step 4: Synthesis of 3-(1-((*3R*, *4R*)-4-(4-(trifluoromethyl)benzyloxy)pyrrolidin-3-yl)-1H-1,2,3-triazol-4-yl)pyridine (Compound 7)

A mixture of compound **6** (4 g, 8.1 mmol) and TFA (5 mL) in DCM (20 mL) was stirred at 0 °C under N_2_ for 2 hours. The mixture was concentrated to leave the crude compound **7** (4 g, crude) as yellow oil, which was used directly for next step. LC-MS (ESI) m/z: 390[M+H]^+^.

#### Step 5: Synthesis of 1-((*3R*, *4R*)-3-(4-(pyridin-3-yl)-1H-1,2,3-triazol-1-yl)-4-(4-(trifluoromethyl)benzyloxy)pyrrolidin-1-yl)prop-2-en-1-one (MYF-03-69)

A mixture of compound **7** (500 mg, 1.28 mmol), acryloyl chloride (115 mg, 1.28 mmol) and TEA (388 mg, 3.84 mmol) in DCM (20 mL) was stirred at rt for 2 hours. The mixture was concentrated and purified by prep-HPLC to obtain **MYF-03-69** as light yellow solid (300 mg, yield 52%). LC-MS (ESI) m/z: 444 [M+H]^+^. ^1^H NMR (400 MHz, CD_3_OD) δ (ppm) 8.91 (d, *J* = 1.8 Hz, 1H), 8.50 (d, *J* = 4.9 Hz, 1H), 8.43 (d, *J* = 4.9 Hz, 1H), 8.18 (dt, *J* = 8.0, 1.8 Hz, 1H), 7.51 (d, *J* = 8.0 Hz, 2H), 7.48-7.38 (m, 3H), 6.60-6.49 (m, 1H), 6.23 (dd, *J* = 16.8, 1.9 Hz, 1H), 5.75-5.67 (m, 1H), 5.41 – 5.30 (m,1H), 4.68 (d, *J* = 4.0 Hz, 2H), 4.60-4.49 (m, 1H), 4.32 – 4.16 (m, 1H), 4.15-3.87 (m, 2H), 3.85 – 3.63 (m, 1H).

**MYF-03-162**

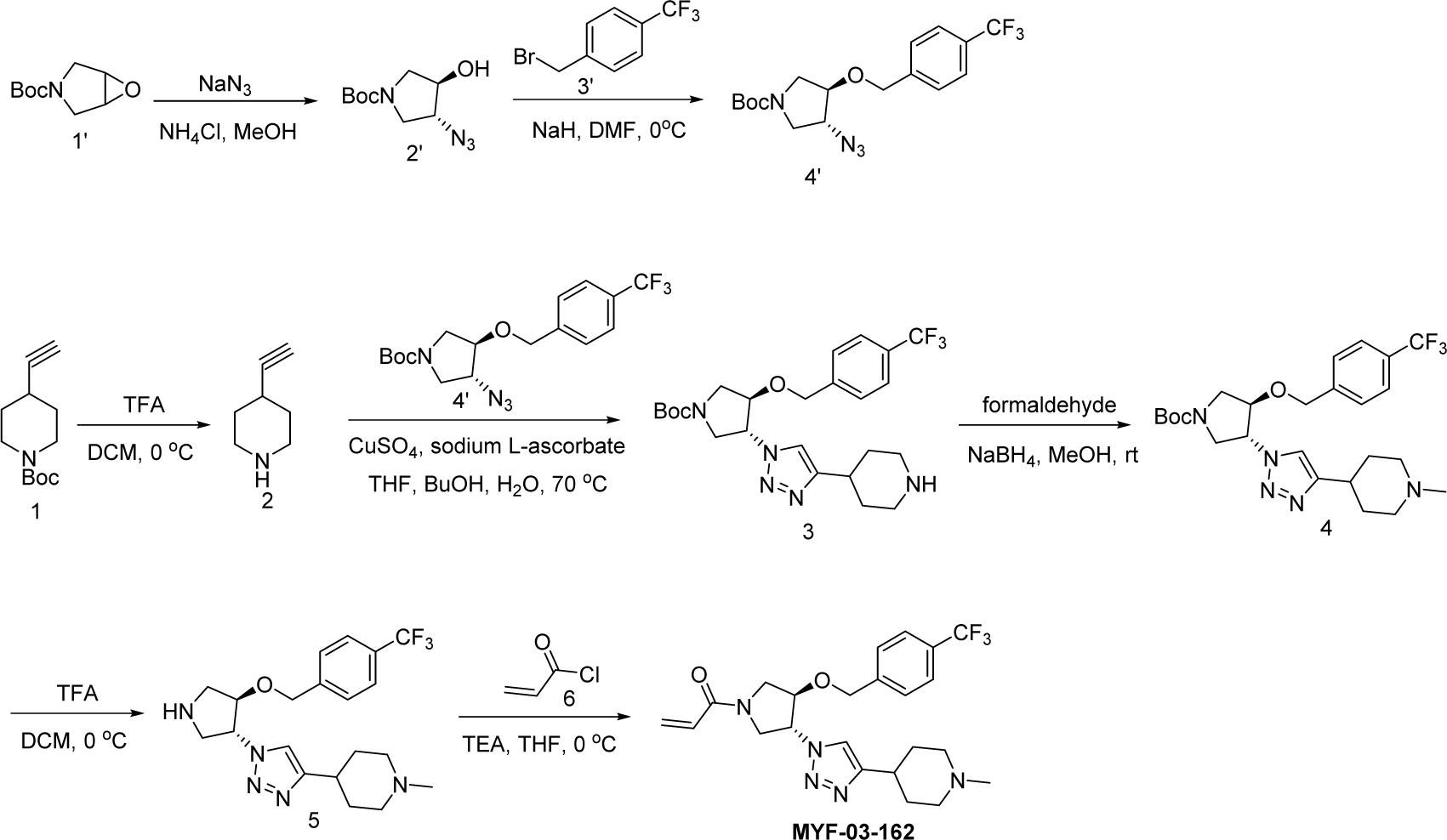

#### Step 1: Synthesis of Compound 2’

A mixture of tert-butyl 6-oxa-3-azabicyclo[3.1.0]hexane-3-carboxylate (10 g, 54 mmol), NaN_3_ (7g, 108 mmol) and NH_4_Cl (2.8g, 54 mmol) in MeOH (120 mL) and H_2_O (20 mL) was stirred at 65°C under N_2_ overnight. The reaction mixture was concentrated in vacuum, the residue was extracted with ethyl acetate (300 mL x 3), the combined organic was washed with water (200 mL x2), dried over anhydrous Na_2_SO_4_, concentrated and purified by flash column chromatography on silica gel (ethyl acetate in petroleum ether = 20% v/v) to obtain compound **2’** (11 g, yield 89%) as oil. LC-MS (ESI) m/z: 129 [M+H-100]^+^.

#### Step 2: Synthesis of Compound 4’

A mixture of compound **2’** (3g, 13.1 mmol), 1-(bromomethyl)-4-(trifluoromethyl)benzene (3.1g, 13.1mmol) and 60% NaH (0.6g, 15.7 mmol) in DMF (20 mL) was stirred at rt under N_2_ protection for 6 hours. The reaction mixture was diluted with water (100 mL) and extracted with ethyl acetate (100 mL x 2), the combined organic was washed with water (100 mL), dried over anhydrous Na_2_SO_4_, concentrated and purified by flash column chromatography on silica gel (ethyl acetate in petroleum ether = 30% v/v) to obtain compound **4’** (3.8g, yield 75%) as oil. LC-MS (ESI) m/z: 287 [M+H-100]^+^.

#### Step 3: Synthesis of 4-ethynylpiperidine (Compound 2)

A mixture of tert-butyl 4-ethynylpiperidine-1-carboxylate (1000 mg, 4.7 mmol) and TFA (0.5 mL) in DCM (1 mL) was stirred at rt under N_2_ protection for 2 hours. The mixture was concentrated to leave the crude compound **2** (1.2 g) as white solid, which was used directly in the next step. LC-MS (ESI) m/z: 110 [M+H]^+^.

#### Step 4: Synthesis of *trans*-tert-butyl 3-(4-(piperidin-4-yl)-1H-1,2,3-triazol-1-yl)-4-(4-(trifluoromethyl)benzyloxy)pyrrolidine-1-carboxylate (Compound 3)

A mixture of compound **4’** (200 mg, 0.52 mmol), compound **2** (113 mg, 1.04 mmol), CuSO_4_ (48 mg, 0.3 mmol) and sodium L-ascorbate (28 mg, 0.14 mmol) in THF (1 mL), H_2_O (1 mL) and ^n^BuOH (1 mL) was stirred at 70 °C under N_2_ overnight. The reaction mixture was concentrated in vacuum, the residue was diluted with water (50 mL) and extracted with ethyl acetate (20 mL x 2), the combined organic was washed with brine (20 mL), dried over anhydrous Na_2_SO_4_, concentrated and purified by flash column chromatography on silica gel (ethyl acetate in petroleum ether = 90% v/v) to obtain compound **3** as yellow solid (200 mg, yield 78%). LC-MS (ESI) m/z: 496[M+H]^+^.

#### Step 5: Synthesis of *trans*-tert-butyl 3-(4-(1-methylpiperidin-4-yl)-1H-1,2,3-triazol-1-yl)-4-(4-(trifluoromethyl)benzyloxy)pyrrolidine-1-carboxylate (Compound 4)

A mixture of compound **3** (100 mg, 0.2 mmol) and formaldehyde (10 mg, 0.3 mmol) and NaBH_4_ (20 mg, 0.4 mmol) in MeOH (10 mL) was stirred at rt for 2 hours. The resulting mixture was concentrated and purified by flash column chromatography on silica gel (ethyl acetate in petroleum ether = 80% v/v) to obtain compound **4** as oil (70 mg, yield 69%). LC-MS (ESI) m/z: 510[M+H]^+^.

#### Step 6: Synthesis of 1-methyl-4-(1-(*trans*-4-(4-(trifluoromethyl)benzyloxy)pyrrolidin-3-yl)-1H-1,2,3-triazol-4-yl)piperidine (Compound 5)

A mixture of compound **4** (70 mg, 0.13 mmol) and TFA (0.5 mL) in DCM (1 mL) was stirred at rt under N_2_ for 2 hours. The mixture was concentrated to leave the crude compound **5** (50 mg) as yellow oil, which was used directly in the next step. LC-MS (ESI) m/z: 410[M+H]^+^.

#### Step 7: Synthesis of 1-(*trans*-3-(4-(1-methylpiperidin-4-yl)-1H-1,2,3-triazol-1-yl)-4-(4-(trifluoromethyl)benzyloxy)pyrrolidin-1-yl)prop-2-en-1-one (Compound MYF-03-162)

To the mixture of compound **5** (50 mg, 0.12 mmol) and TEA (30 mg, 0.24 mmol) in DCM (10 mL) was added acryloyl chloride (20 mg, 0.12 mmol), the mixture was stirred at rt under N_2_ for 2 hours, and then concentrated and purified by prep-HPLC to obtain compound **MYF-03-162** as white solid (13 mg, yield 23%). LC-MS (ESI) m/z: 464 [M+H]^+^.; ^1^H NMR (400 MHz, CD_3_OD) δ (ppm) 7.89 (d, *J* = 3.6 Hz, 1H), 7.63 (d, *J* = 8.1 Hz, 2H), 7.49(d, *J* = 8.1 Hz, 2H), 6.61 (dd, *J* = 16.8, 10.4 Hz, 1H), 6.31 (dd, *J* = 16.8, 1.8 Hz, 1H), 5.78 (ddd, *J* = 10.4, 3.6, 1.9 Hz, 1H), 5.39 – 5.27 (m, 1H), 4.73 (d, *J* = 4.6 Hz, 2H), 4.61-4.50 (m, 1H), 4.34 – 4.10 (m, 2H), 4.09-3.69 (m, 2H), 2.94 (d, *J* = 11.6 Hz, 2H), 2.80-2.68 (m, 1H), 2.31 (s, 3H), 2.24 – 2.12 (m, 2H), 2.08 – 1.98 (m, 2H), 1.80-1.65 (m, 2H).

**MYF-03-139**

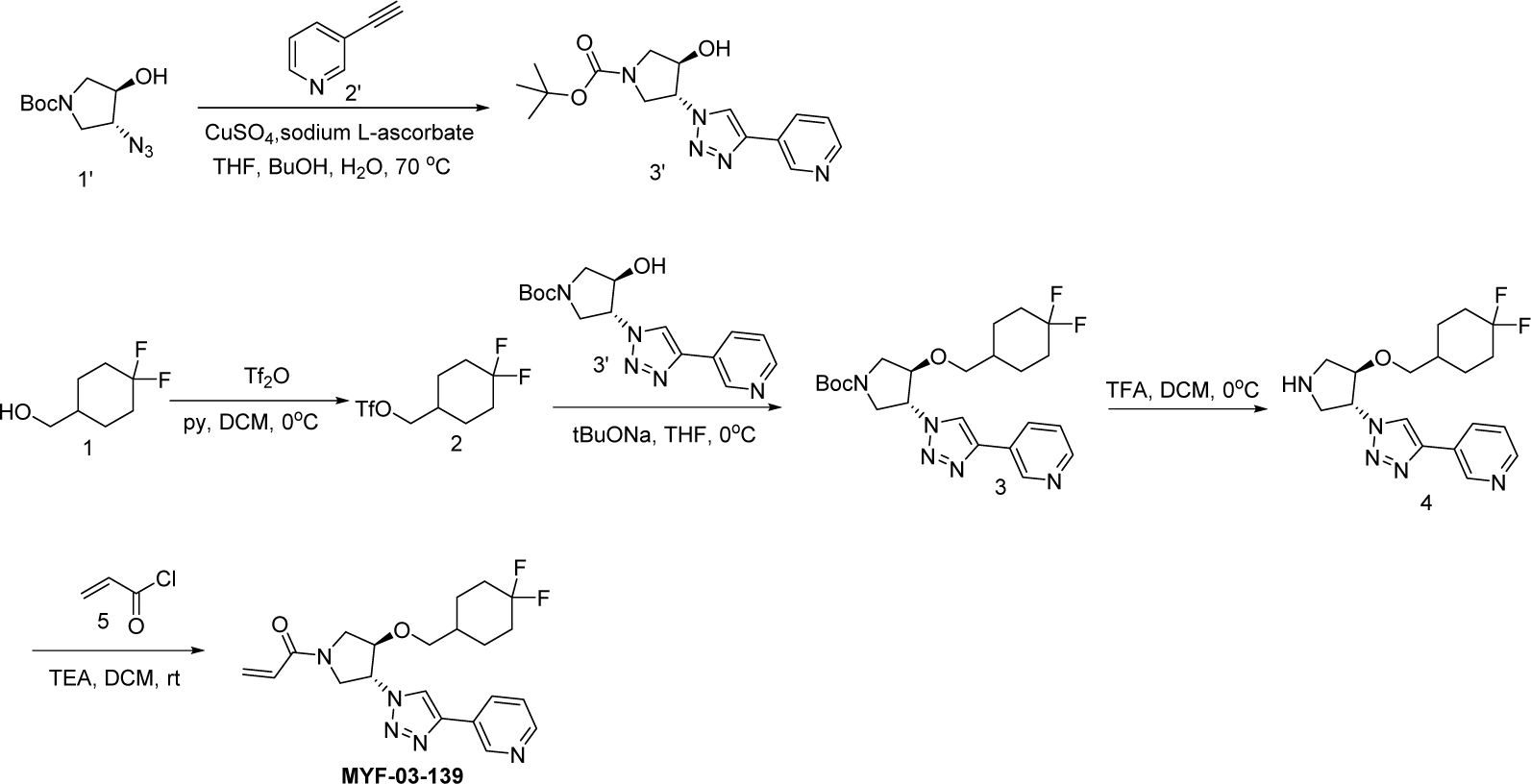

#### Step 1: Synthesis of *trans*-tert-butyl 3-hydroxy-4-(4-(pyridin-3-yl)-1H-1,2,3-triazol-1-yl)pyrrolidine-1-carboxylate (Compound 3’)

A mixture of tert-butyl compound **1’** (1 g, 4.38 mmol), 3-ethynylpyridine (451 mg, 4.38 mmol), CuSO_4_ (654 mg, 2.6 mmol) and sodium L-ascorbate (257 mg, 1.3 mmol) in THF (3 mL), H_2_O (3 mL) and ^n^BuOH (3 mL) was stirred at 70 °C under N_2_ protection overnight. The reaction mixture was diluted with ethyl acetate (60mL), washed with water (40 mL), dried over anhydrous Na_2_SO_4_, concentrated and purified by flash column chromatography on silica gel (ethyl acetate in petroleum ether = 80% v/v) to obtain compound **3’** (650 mg, yield 45%) as yellow solid. LC-MS (ESI) m/z: 332[M+H]^+^.

#### Step 2: Synthesis of (4,4-difluorocyclohexyl)methyl trifluoromethanesulfonate (Compound 2)

A mixture of (4,4-difluorocyclohexyl)methanol (1 g, 6.6 mmol), Tf_2_O (2.81 g, 9.9 mmol) and pyridine (1 mL) in DCM (20 mL) was stirred at rt under N_2_ protection for 3 hours. The reaction mixture was diluted with ethyl acetate (100 mL), washed with water (100 mL x 2), dried over anhydrous Na_2_SO_4_, filtered and concentrated to leave the crude compound **2** (500 mg) as yellow oil, which was used directly in the next step. LC-MS (ESI) m/z: no MS.

#### Step 3: Synthesis of *trans*-tert-butyl 3-((4,4-difluorocyclohexyl)methoxy)-4-(4-(pyridin-3-yl)-1H-1,2,3-triazol-1-yl)pyrrolidine-1-carboxylate (Compound 3)

A mixture of compound **2** (200 mg, 0.7 mmol), compound **3’** (58 mg, 0.175 mmol) and ^t^BuONa (25 mg, 0.26 mmol) in THF (5 mL) was stirred at 0 °C under N_2_ for 2 hours. The reaction mixture was diluted with ethyl acetate (50 mL), washed with water (50 mL x 2), dried over anhydrous Na_2_SO_4_, concentrated and purified by flash column chromatography on silica gel (ethyl acetate in petroleum ether = 50% v/v) to obtain compound **3** as oil (100 mg, yield 30%). LC-MS (ESI) m/z: 464[M+H]^+^.

#### Step 4: Synthesis of 3-(1-(*trans*-4-((4,4-difluorocyclohexyl)methoxy)pyrrolidin-3-yl)-1H-1,2,3-triazol-4-yl)pyridine (Compound 4)

A mixture of compound **3** (50 mg, 0.1 mmol) and TFA (1 mL) in DCM (3 mL) was stirred at rt for 2 hours. The mixture was concentrated to leave the crude compound **4** (50 mg) as yellow oil, which was used directly in the next step. LC-MS (ESI) m/z: 364[M+H]^+^.

#### Step 5: Synthesis of 1-(*trans*-3-((4,4-difluorocyclohexyl)methoxy)-4-(4-(pyridin-3-yl)-1H-1,2,3-triazol-1-yl)pyrrolidin-1-yl)prop-2-en-1-one (Compound MYF-03-139)

A mixture of compound **4** (50 mg, 0.1 mmol), acryloyl chloride (10 mg, 0.1 mmol), and TEA (20 mg, 0.2 mmol) in DCM (5 mL) was stirred at rt under N_2_ for 2 hours. The mixture was diluted with DCM (50 mL), washed with water (50 mL x 2), dried over anhydrous Na_2_SO_4_, concentrated and purified by prep-HPLC to obtain **MYF-03-139** as white solid (16 mg, yield 38%). LC-MS (ESI) m/z: 418[M+H]^+^; ^1^H-NMR (400 MHz, CD_3_OD) δ (ppm) 8.91 (d, *J* = 1.2 Hz, 1H), 8.55 – 8.40 (m, 2H), 8.18 (dt, *J* = 8.0, 1.9 Hz, 1H), 7.43 (dd, *J* = 8.0, 4.9 Hz, 1H), 6.55 (ddd, *J* = 16.8, 10.4, 2.5 Hz, 1H), 6.28 – 6.20 (m, 1H), 5.70 (ddd, *J* = 10.5, 4.7, 1.9 Hz, 1H), 5.29 – 5.18 (m, 1H), 4.43 – 4.32 (m, 1H), 4.27 – 4.16 (m, 1H), 4.07 – 3.37 (m, 5H), 1.91 (ddd, *J* = 13.9, 7.0, 3.5 Hz, 2H), 1.76 – 1.54 (m, 5H), 1.26 – 1.13 (m, 2H).

**MYF-03-137**

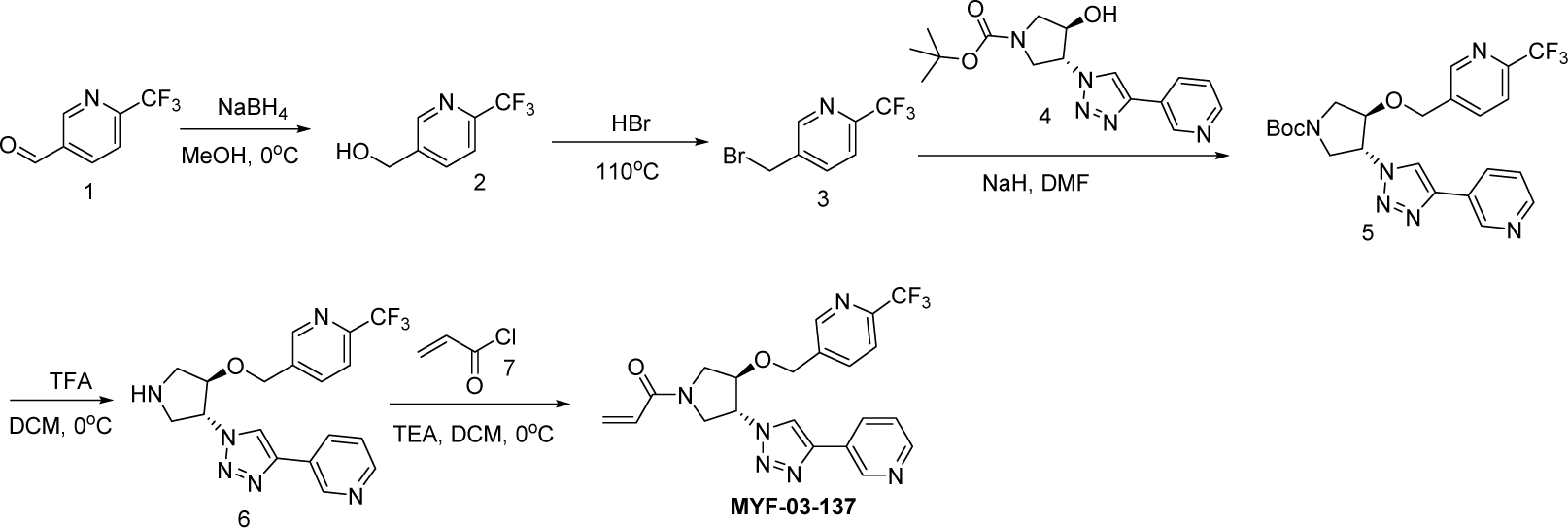

#### Step 1: Synthesis of (6-(trifluoromethyl)pyridin-3-yl)methanol (Compound 2)

A mixture of 6-(trifluoromethyl)nicotinaldehyde (170 mg, 1 mmol) and NaBH_4_ (76 mg, 2 mmol) in MeOH (3 mL) was stirred at 0 °C for 3 hours. The reaction mixture was concentrated in vacuum, the residue was extracted with ethyl acetate (60 mL), washed with water (40 mL), dried over anhydrous Na_2_SO_4_, filtered and concentrated to leave the crude product (150 mg, yield 84%) as oil, which was used directly in the next step. LC-MS (ESI) m/z: 178[M+H]^+^.

#### Step 2: Synthesis of 5-(bromomethyl)-2-(trifluoromethyl)pyridine (Compound 3)

A mixture of compound **2** (400 mg, 2.2 mmol) and 48% aqueous HBr solution (6 mL) was stirred at 110 °C overnight. The reaction mixture was concentrated in vacuum, the residue was diluted with ethyl acetate (60 mL), washed with water (40 mL), dried over anhydrous Na_2_SO_4_, concentrated and purified by flash column chromatography on silica gel (ethyl acetate in petroleum ether = 20% v/v) to obtain compound **3** as oil (200 mg, yield 38%). LC-MS (ESI) m/z: 240[M+H]^+^.

#### Step 3: Synthesis of *trans*-tert-butyl 3-(4-(pyridin-3-yl)-1H-1,2,3-triazol-1-yl)-4-((6-(trifluoromethyl)pyridin-3-yl)methoxy)pyrrolidine-1-carboxylate (Compound 5)

A mixture of compound **3** (100 mg, 0.4 mmol), compound **4** (140 mg, 0.4 mmol) and 60% NaH (32 mg, 0.8 mmol) in DMF (5 mL) was stirred at rt under N_2_ overnight. The reaction mixture was diluted with water (100 mL) and extracted with ethyl acetate (20 mL x 2), the combined organic was washed with water (50 mL), dried over anhydrous Na_2_SO_4_, concentrated and purified by flash column chromatography on silica gel (ethyl acetate in petroleum ether = 90% v/v) to obtain compound **5** as solid (100 mg, yield 51%). LC-MS (ESI) m/z: 491[M+H]^+^.

#### Step 4: Synthesis of 5-((*trans*-4-(4-(pyridin-3-yl)-1H-1,2,3-triazol-1-yl)pyrrolidin-3-yloxy)methyl)-2-(trifluoromethyl)pyridine (Compound 6)

A mixture of compound **5** (100 mg, 0.2 mmol) and TFA (1 mL) in DCM (3 mL) was stirred at rt for 2 hours. The mixture was concentrated to leave the crude compound **6** (100 mg, crude) as yellow oil, which was used directly in the next step. LC-MS (ESI) m/z: 391

#### Step 5: Synthesis of 1-(*trans*-3-(4-(pyridin-3-yl)-1H-1,2,3-triazol-1-yl)-4-((6-(trifluoromethyl)pyridin-3-yl)methoxy)pyrrolidin-1-yl)prop-2-en-1-one (Compound MYF-03-137)

A mixture of compound **6** (50 mg, 0.125 mmol), acryloyl chloride (15 mg, 0.125 mmol) and TEA (25 mg, 0.25 mmol) in DCM (3 mL) was stirred at 0 °C for 2 hours. The mixture was concentrated and purified by prep-HPLC to obtain **MYF-03-137** as white solid (5 mg, yield 9%). LC-MS (ESI) m/z: 445. ^1^H NMR (400 MHz, CD_3_OD) δ (ppm) 9.04 (d, *J* = 2.2 Hz, 1H), 8.76 – 8.50 (m, 3H), 8.31 (dt, *J* = 7.9, 1.9 Hz, 1H), 8.03 (d, *J* = 8.2 Hz, 1H), 7.80 (d, *J* = 8.1 Hz, 1H), 7.56 (dd, *J* = 8.0, 4.9 Hz, 1H), 6.67 (ddd, *J* = 16.8, 10.4, 3.5 Hz, 1H), 6.36 (dd, *J* = 16.8, 1.7 Hz, 1H), 5.83 (ddd, *J* = 10.5, 5.1, 1.9 Hz, 1H), 5.58 – 5.44 (m, 1H), 4.97 (s, 2H), 4.81 – 4.68 (m, 1H), 4.48 – 4.03 (m, 3H), 4.02 – 3.80 (m, 1H).

**MYF-03-138**

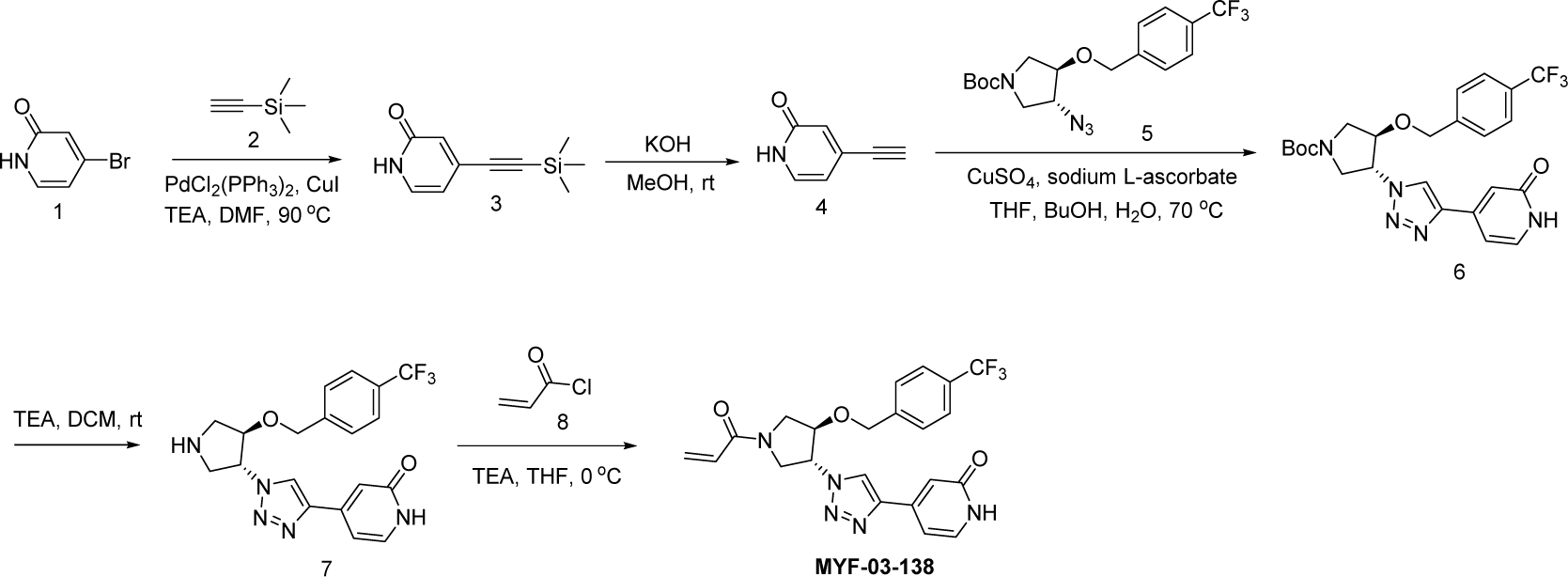

#### Step 1: Synthesis of 4-((trimethylsilyl)ethynyl)pyridin-2(1H)-one (Compound 3)

To the solution of ethynyltrimethylsilane (0.595 g, 6.07 mmol) in DMF (40 mL) was added 4-bromopyridin-2(1H)-one (1 g, 5.8 mmol), PdCl_2_(PPh_3_)_2_ (0.204 g, 0.29 mmol), CuI (55 mg, 0.29 mmol) and Et_3_N (1.17 g, 11.6 mmol). The mixture was stirred at 90 °C under N_2_ for 2 hours. After cooled down to rt the mixture was diluted with water (200 mL) and extracted with EtOAc (50 mL x 2), the combined organic was dried over anhydrous Na_2_SO_4_, concentrated and purified by flash column chromatography on silica gel (ethyl acetate in petroleum ether = 50% v/v) to obtain compound **3** as oil (500 mg, yield 45.4%). LC-MS (ESI) m/z: 192[M+H]^+^.

#### Step 2: Synthesis of 4-ethynylpyridin-2(1H)-one (Compound 4)

To the solution of compound **3** (450 mg, 2.35 mmol) in MeOH (20 mL) was added KOH (263 mg, 4.70 mmol). The mixture was stirred at rt under N_2_ for 2 hours. The resulted mixture was concentrated and purified by flash column chromatography on silica gel (ethyl acetate in petroleum ether = 90% v/v) to obtain compound **4** as oil (200 mg, yield 71.4%). LC-MS (ESI) m/z: 120[M+H]^+^.

#### Step 3: Synthesis of *trans*-tert-butyl 3-(4-(2-oxo-1,2-dihydropyridin-4-yl)-1H-1,2,3-triazol-1-yl)-4-(4-(trifluoromethyl)benzyloxy)pyrrolidine-1-carboxylate (Compound 6)

To the solution of compound **5** (250 mg, 0.65 mmol) in THF (10mL), H_2_O (10mL) and ^n^BuOH (10mL) was added compound **4** (116 mg, 0.97 mmol), CuSO_4_ (15 mg, 0.065 mmol) and sodium L-ascorbate (26 mg, 0.13 mmol). The mixture was stirred at 70 °C under N_2_ for 16 hours. The resulting mixture was concentrated and purified by flash column chromatography on silica gel (methanol in dichloromethane = 20% v/v) to obtain compound **6** as solid (200 mg, yield 60%). LC-MS (ESI) m/z: 506[M+H]+.

#### Step 4: Synthesis of 4-(1-(*trans*-4-(4-(trifluoromethyl)benzyloxy)pyrrolidin-3-yl)-1H-1,2,3-triazol-4-yl)pyridin-2(1H)-one (Compound 7)

To the solution of compound **6** (180 mg, 0.36 mmol) in DCM (10 mL) was added TFA (2 mL). The mixture was stirred at rt for 2 hours and concentrated in vacuum, the residue was adjusted to pH∼8 with NaHCO_3_ solution and extracted with EtOAc (50 mL x 3), the combined organics were washed with brine (100 mL), dried over anhydrous Na_2_SO_4_, filtered and concentrated to leave crude compound **7** as oil (150 mg, crude). LC-MS (ESI) m/z: 406[M+H]^+^.

#### Step 5: Synthesis of 4-(1-(*trans*-1-acryloyl-4-(4-(trifluoromethyl)benzyloxy)pyrrolidin-3-yl)-1H-1,2,3-triazol-4-yl)pyridin-2(1H)-one (Compound MYF-03-138)

To the solution of compound **7** (130 mg, 0.32 mmol) in THF (10 mL) was added acryloyl chloride (29 mg, 0.32 mmol) and Et_3_N (65 mg, 0.64 mmol). The mixture was stirred at 0°C for 1 hour, and then concentrated and purified by prep-HPLC to obtain compound **MYF-03-138** as solid (44 mg, yield 30.0%). LC-MS (ESI) m/z: 460[M+H]^+^. ^1^H-NMR (400 MHz, DMSO-*d_6_*) δ (ppm) 11.58 (br, 1H), 8.89 (d, *J* = 5.5 Hz, 1H), 7.70 (d, *J* = 8.0 Hz, 2H), 7.54 (d, *J* = 8.0 Hz, 2H), 7.46 (d, *J* = 6.6 Hz, 1H), 6.79 (s, 1H), 6.70 – 6.56 (m, 2H), 6.24 – 6.14 (m, 1H), 5.79-5.71 (m, 1H), 5.53 – 5.39 (m, 1H), 4.81 – 4.71 (m, 2H), 4.62-4.50 (m, 1H), 4.30-4.13 (m, 1H), 4.12-3.95 (m, 1H), 3.91-3.79 (m, 1H), 3.68-3.58 (m, 1H).

**MYF-03-146**

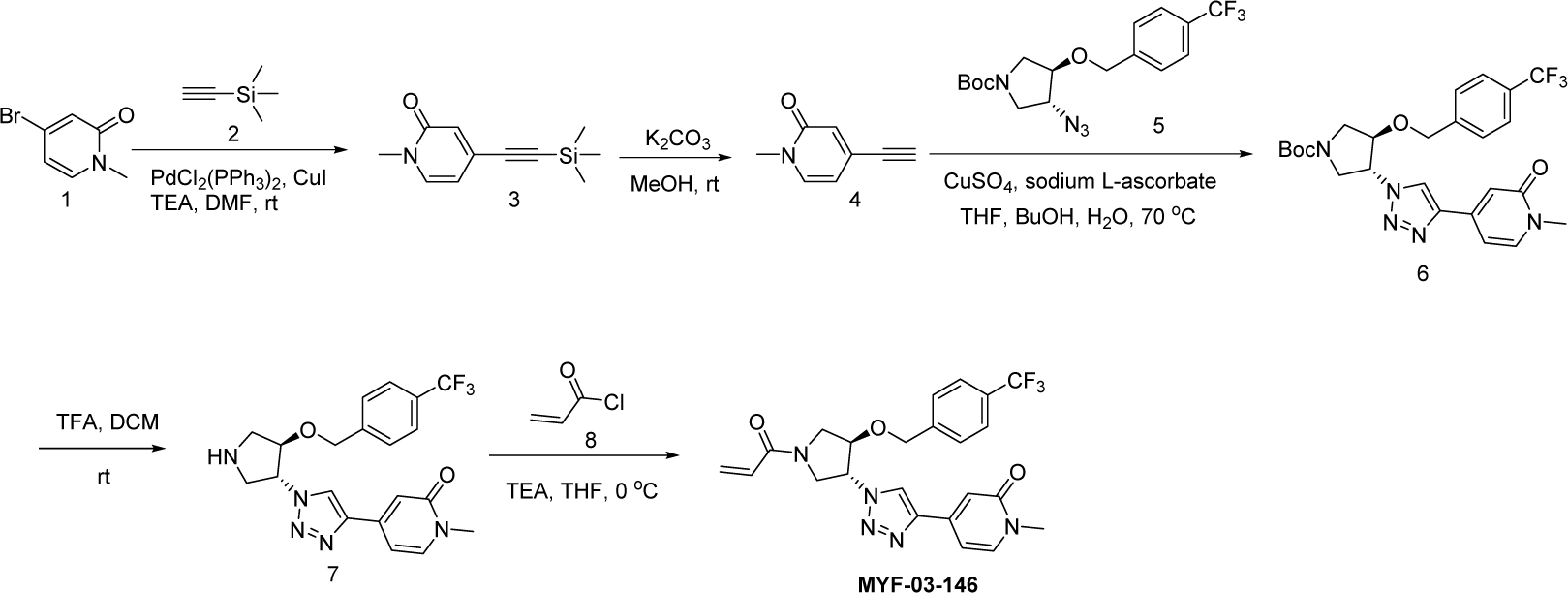

#### Step 1: Synthesis of 1-methyl-4-((trimethylsilyl)ethynyl)pyridin-2(1H)-one (Compound 3)

To the solution of 4-bromo-1-methylpyridin-2(1H)-one (1 g, 5.3 mmol) in DMF (20 mL) was added ethynyltrimethylsilane (0.55 g, 5.6 mmol), PdCl_2_(PPh_3_)_2_ (0.21 g, 0.3 mmol), CuI (0.06 g, 0.3 mmol) and Et_3_N (1.07 g, 10.6 mmol). The mixture was stirred at room temperature under N_2_ for 2 hours. The resulting mixture was concentrated and purified by flash column chromatography on silica gel (ethyl acetate in petroleum ether = 50% v/v) to obtain compound **3** as oil (1.1 g, yield 99.9%). LC-MS (ESI) m/z: 206[M+H]^+^.

#### Step 2: Synthesis of 4-ethynyl-1-methylpyridin-2(1H)-one (Compound 4)

To the solution of compound **3** (1 g, 4.9 mmol) in MeOH (20 mL) was added K_2_CO_3_ (1.35 g, 9.8 mmol). The mixture was stirred at room temperature under N_2_ for 2 hours. The resulting mixture was concentrated and purified by flash column chromatography on silica gel (ethyl acetate in petroleum ether = 90% v/v) to obtain compound **4** as solid (400 mg, yield 56.1%). LC-MS (ESI) m/z: 134[M+H]^+^.

#### Step 3: Synthesis of *trans*-tert-butyl 3-(4-(1-methyl-2-oxo-1,2-dihydropyridin-4-yl)-1H-1,2,3-triazol-1-yl)-4-(4-(trifluoromethyl)benzyloxy)pyrrolidine-1-carboxylate (Compound 6)

To the solution of *trans*-tert-butyl 3-azido-4-(4-(trifluoromethyl)benzyloxy) pyrrolidine-1-carboxylate (350 mg, 0.91 mmol) in THF (5mL), H_2_O (5mL) and ^n^BuOH (5mL) was added compound **4** (181 mg, 1.36 mmol), CuSO_4_ (23 mg, 0.09 mmol) and sodium L-ascorbate (36 mg, 0.18 mmol). The mixture was stirred at 70 °C under N_2_ for 16 hours. The resulting mixture was concentrated and purified by flash column chromatography on silica gel (methanol in dichloromethane=20% v/v) to obtain compound **6** as solid (300 mg, yield 63.5%). LC-MS (ESI) m/z: 520[M+H]^+^.

#### Step 4: Synthesis of 1-methyl-4-(1-(*trans*-4-(4-(trifluoromethyl)benzyloxy)pyrrolidin-3-yl)-1H-1,2,3-triazol-4-yl)pyridin-2(1H)-one (Compound 7)

To the solution of compound **6** (280 mg, 0.54 mmol) in DCM (10 mL) was added TFA (2 mL). The mixture was stirred at room temperature for 2 hours and concentrated in vacuum, the residue was adjusted to pH∼8 with NaHCO_3_ solution and extracted with EtOAc (50 mL x 3), the combined organics were washed with brine (100 mL), dried over anhydrous Na_2_SO_4_, filtered and concentrated to leave the crude compound **7** as oil (200 mg, crude). LC-MS (ESI) m/z: 420[M+H]^+^.

#### Step 5: Synthesis of 4-(1-(*trans*-1-acryloyl-4-(4-(trifluoromethyl)benzyloxy)pyrrolidin-3-yl)-1H-1,2,3-triazol-4-yl)-1-methylpyridin-2(1H)-one (Compound MYF-03-146)

To the solution of compound **7** (180 mg, 0.43 mmol) in THF (10 mL) was added acryloyl chloride (39 mg, 0.43 mmol) and Et_3_N (87 mg, 0.86 mmol). The mixture was stirred at 0°C under N_2_ for 1 hour, and then concentrated and purified by prep-HPLC to obtain compound **MYF-03-146** as solid (215 mg, yield 95.5%). LC-MS (ESI) m/z: 474[M+H]^+^. ^1^H-NMR (400 MHz, DMSO-*d_6_*): δ (ppm) 8.89 (d, *J* = 5.8 Hz, 1H), 7.78 (d, *J* = 7.1 Hz, 1H), 7.70 (d, *J* = 8.2 Hz, 2H), 7.53 (d, *J* = 8.1 Hz, 2H), 6.85 (s, 1H), 6.76 – 6.53 (m, 2H), 6.19 (dd, *J* = 16.8, 2.3 Hz, 1H), 5.73 (ddd, *J* = 10.2, 5.1, 2.3 Hz, 1H), 5.47 (d, *J* = 25.4 Hz, 1H), 4.75 (s, 2H), 4.57 (d, *J* = 24.5 Hz, 1H), 4.31 – 3.57 (m, 4H), 3.44 (s, 3H).

**MYF-03-135**

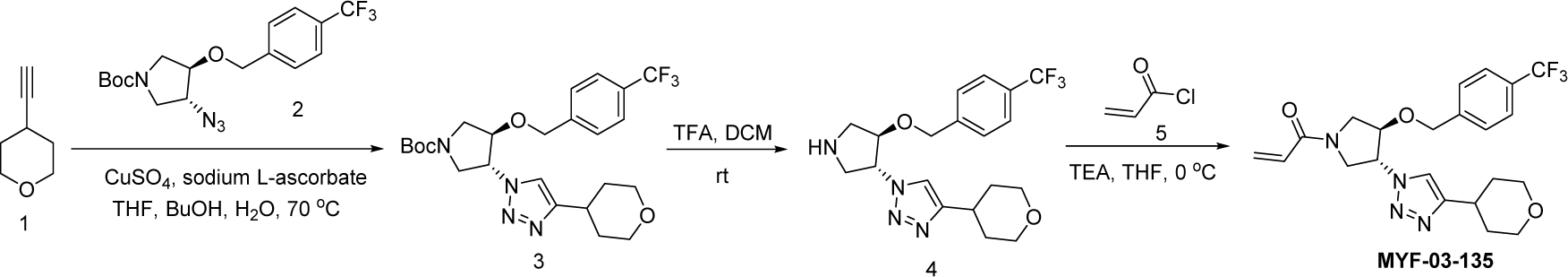

#### Step 1: Synthesis of *trans*-tert-butyl 3-(4-(tetrahydro-2H-pyran-4-yl)-1H-1,2,3-triazol-1-yl)-4-(4-(trifluoromethyl)benzyloxy)pyrrolidine-1-carboxylate (Compound 3)

To the solution of compound **2** (300 mg, 0.78 mmol) in THF (10 mL), H_2_O (10 mL) and ^n^BuOH (10 mL) was added 4-ethynyltetrahydro-2H-pyran (128 mg, 1.16 mmol), CuSO_4_ (20 mg, 0.078 mmol) and sodium L-ascorbate (31 mg, 0.156 mmol). The mixture was stirred at 70 °C under N_2_ for 16 hours. The resulting mixture was concentrated and purified by flash column chromatography on silica gel (Methanol in dichloromethane = 20% v/v) to obtain compound **3** as solid (300 mg, yield 77.9%). LC-MS (ESI) m/z: 497 [M+H]^+^.

#### Step 2: Synthesis of 4-(tetrahydro-2H-pyran-4-yl)-1-(*trans*-4-(4-(trifluoromethyl)benzyloxy)pyrrolidin-3-yl)-1H-1,2,3-triazole (Compound 4)

To the solution of compound **3** (280 mg, 0.56 mmol) in DCM (10 mL) was added TFA (2 mL). The mixture was stirred at room temperature for 2 hours and concentrated in vacuum, the residue was adjusted to pH∼8 with NaHCO_3_ solution and extracted with EtOAc (50 mL x 3), the combined organics were washed with brine (100 mL), dried over anhydrous Na_2_SO_4_, filtered and concentrated to leave the crude compound **4** as oil (250 mg, crude). LC-MS (ESI) m/z: 397 [M+H]^+^.

#### Step 3: Synthesis of 1-(*trans*-3-(4-(tetrahydro-2H-pyran-4-yl)-1H-1,2,3-triazol-1-yl)-4-(4-(trifluoromethyl)benzyloxy)pyrrolidin-1-yl)prop-2-en-1-one (Compound MYF-03-135)

To the solution of compound **4** (230 mg, 0.58 mmol) in THF (10 mL) was added acryloyl chloride (52 mg, 0.58 mmol) and Et_3_N (117 mg, 1.16 mmol). The mixture was stirred at 0°C under N_2_ for 1 hour, and then concentrated and purified by prep-HPLC to obtain compound **MYF-03-135** as solid (180 mg, yield 69.0%). LC-MS (ESI) m/z: 451[M+H]^+^. ^1^H-NMR (400 MHz, DMSO-*d_6_*): δ (ppm) 8.05 (d, *J* = 6.6 Hz, 1H), 7.70 (d, *J* = 7.6 Hz, 2H), 7.51 (d, *J* = 7.8 Hz, 2H), 6.66 – 6.57 (m, 1H), 6.18 (d, *J* =16.8 Hz, 1H), 5.77 – 5.67 (m, 1H), 5.43 – 5.29 (m, 1H), 4.72 (d, *J* = 5.2 Hz, 2H), 4.50 (dd, *J* = 31.4, 5.0 Hz, 1H), 4.24 – 3.96 (m, 2H), 3.89 (d, *J* = 13.1 Hz, 2H), 3.82 (td, *J* = 13.1, 4.4 Hz, 1H), 3.58 (dd, *J* = 13.1, 3.3 Hz, 1H), 3.44 (t, *J* = 11.3 Hz, 2H), 2.93 (t, *J* = 10.6 Hz, 1H), 1.85 (d, *J* = 12.9 Hz, 2H), 1.61 (q, *J* = 11.4 Hz, 2H).

**MYF-03-176**

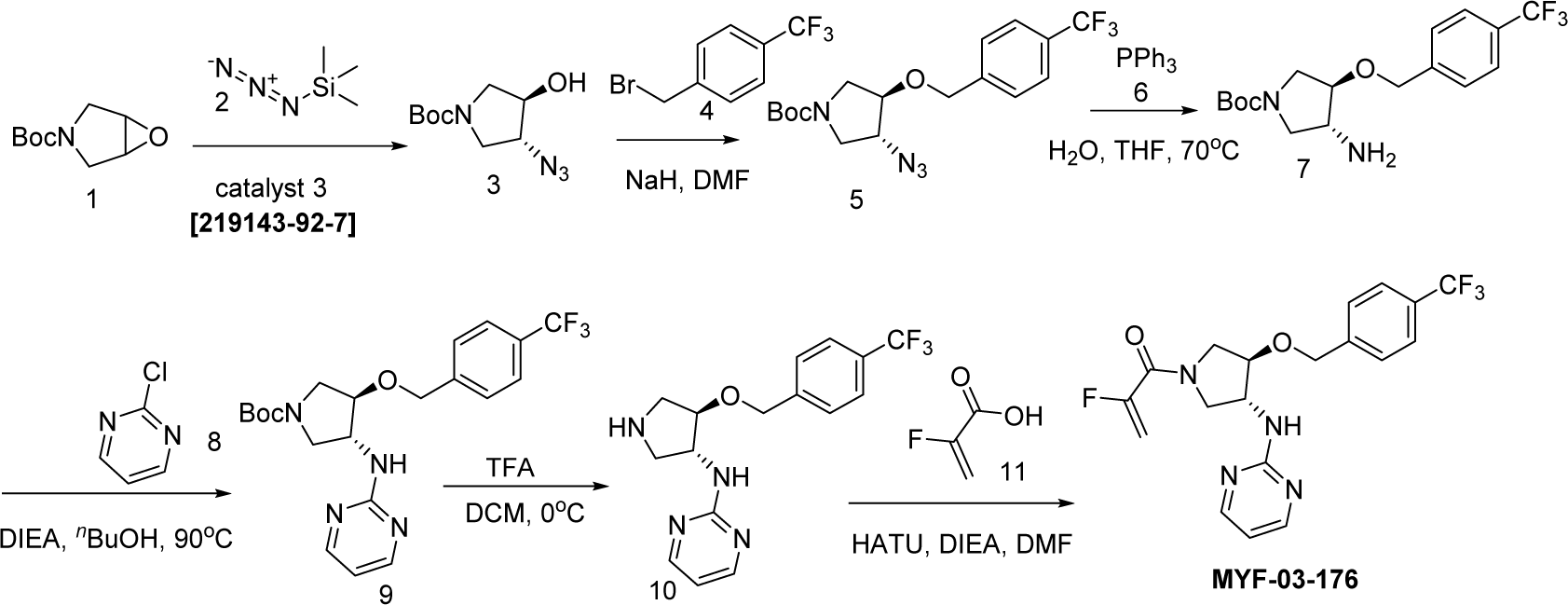

#### Step 1: Synthesis of (3R,4R)-tert-butyl 3-azido-4-hydroxypyrrolidine-1-carboxylate (Compound 3)

A mixture of compound **1** (4000mg, 21.6mmol), compound **2** (2664mg, 23.2mmol) and catalyst **3** (328mg, 0.42mmol) was stirred at rt for overnight under N_2_ protection. The reaction mixture was treated with MeOH (60mL) and K_2_CO_3_ (1788mg, 12.8mmol) and continued to stirring for 5h. The reaction mixture was extracted with ethyl acetate (300mL x 3), and washed by water (300 mL x 2). The organic layer was dried over Na_2_SO_4_ and concentrated. The residue was purified by flash column chromatography on silica gel (ethyl acetate in petroleum ether = 20% v/v) to obtain 3.5 g product compound **3** as yellow oil (3.5g, yield 71%). LC-MS (ESI) m/z: 129[M+H]^+^.

#### Step 2: Synthesis of (3R,4R)-tert-butyl 3-azido-4-(4-(trifluoromethyl)benzyloxy)pyrrolidine-1-carboxylate (Compound 5)

A mixture of compound **3** (500mg, 2.19 mmol), compound **4** (524mg, 2.19mmol) and NaH (105mg, 2.62 mmol) in THF (10 mL) was stirred at rt for 6h under N_2_ protection. The reaction mixture was monitored by LC-MS. The reaction mixture was extracted with ethyl acetate (100mL), and washed by water (50mL). The organic layer was dried over Na_2_SO_4_, was concentrated and purified by column chromatography on silica gel (ethyl acetate in petroleum ether = 30% v/v) to obtain 600mg product compound **5** as white oil (600mg, yield 95%). LC-MS (ESI) m/z: 287[M+ H]^+^.

#### Step 3: Synthesis of (3R,4R)-tert-butyl 3-amino-4-(4-(trifluoromethyl)benzyloxy)pyrrolidine-1-carboxylate (Compound 7)

A mixture of compound **5** (1000mg, 2.58 mmol), PPh_3_ (814mg, 3.1mmol) and H_2_O (930mg, 51.6 mmol) in THF (40 mL) was stirred at 70 °C for 5h under N_2_ protection. The reaction mixture was monitored by LC-MS. The reaction mixture was extracted with ethyl acetate (300mL), and washed by water (200mL), The organic layer was dried over Na_2_SO_4_, concentrated and purified by p-HPLC to obtain 800mg product compound **7** as yellow oil (800mg, yield 86%). LC-MS (ESI) m/z: 261[M+H-100]^+^.

#### Step 4: Synthesis of (3R,4R)-tert-butyl 3-(pyrimidin-2-ylamino)-4-(4-(trifluoromethyl)benzyloxy)pyrrolidine-1-carboxylate (Compound 9)

A mixture of compound **7** (600 mg, 1.6mmol), compound **8** (240 mg, 1.84mmol) and DIPEA (420mg, 3.24 mmol) in *n*BuOH (6 mL) was stirred at 70 °C for overnight under N_2_ protection. The reaction mixture was monitored by LC-MS. The reaction mixture was concentrated and purified by p-HPLC to obtain 500mg product compound **9** as white oil (500mg, yield 71%). LC-MS (ESI) m/z: 439[M+H]^+^.

#### Step 5: Synthesis of N-((3R,4R)-4-(4-(trifluoromethyl)benzyloxy)pyrrolidin-3-yl)pyrimidin-2-amine (Compound 10)

A mixture of compound **9** (400mg, 0.91 mmol) and TFA (1mL) in DCM (3 mL) was stirred at rt for 2h under N_2_ protection. The mixture was concentrated to obtain 300mg crude of compound **10** as yellow oil. Used for next step (400mg, yield 97%). LC-MS (ESI) m/z: 339[M+H]^+^.

#### Step 6: Synthesis of 2-fluoro-1-((3R,4R)-3-(pyrimidin-2-ylamino)-4-(4-(trifluoromethyl)benzyloxy)pyrrolidin-1-yl)prop-2-en-1-one (MYF-03-176)

A mixture of compound **10** (200mg, 0.58 mmol), compound **11** (60mg, 0.69mmol) and HATU (256mg, 0.69 mmol) and DIEA (224mg, 1.74mmol) in DMF (5 mL) was stirred rt for overnight under N_2_ protection. The reaction mixture was monitored by LC-MS. The reaction mixture was concentrated and purified by p-HPLC to obtain 150mg product **MYF-03-176** as white solid (150mg, yield 63%). LC-MS (ESI) m/z: 411[M+H]^+^. 1H NMR (400 MHz, MeOD) δ 8.44 (s, 2H), 7.82 – 7.50 (m, 4H), 6.83 (dd, J = 8.3, 4.9 Hz, 1H), 5.50 (dt, J = 47.1, 3.3 Hz, 1H), 5.27 (dt, J = 16.5, 3.4 Hz, 1H), 4.82 (dd, J = 13.3, 8.8 Hz, 2H), 4.68 – 4.57 (m, 1H), 4.28 – 3.67 (m, 5H)

## SUPPLEMENTARY FIGURES

**Figure S1.**
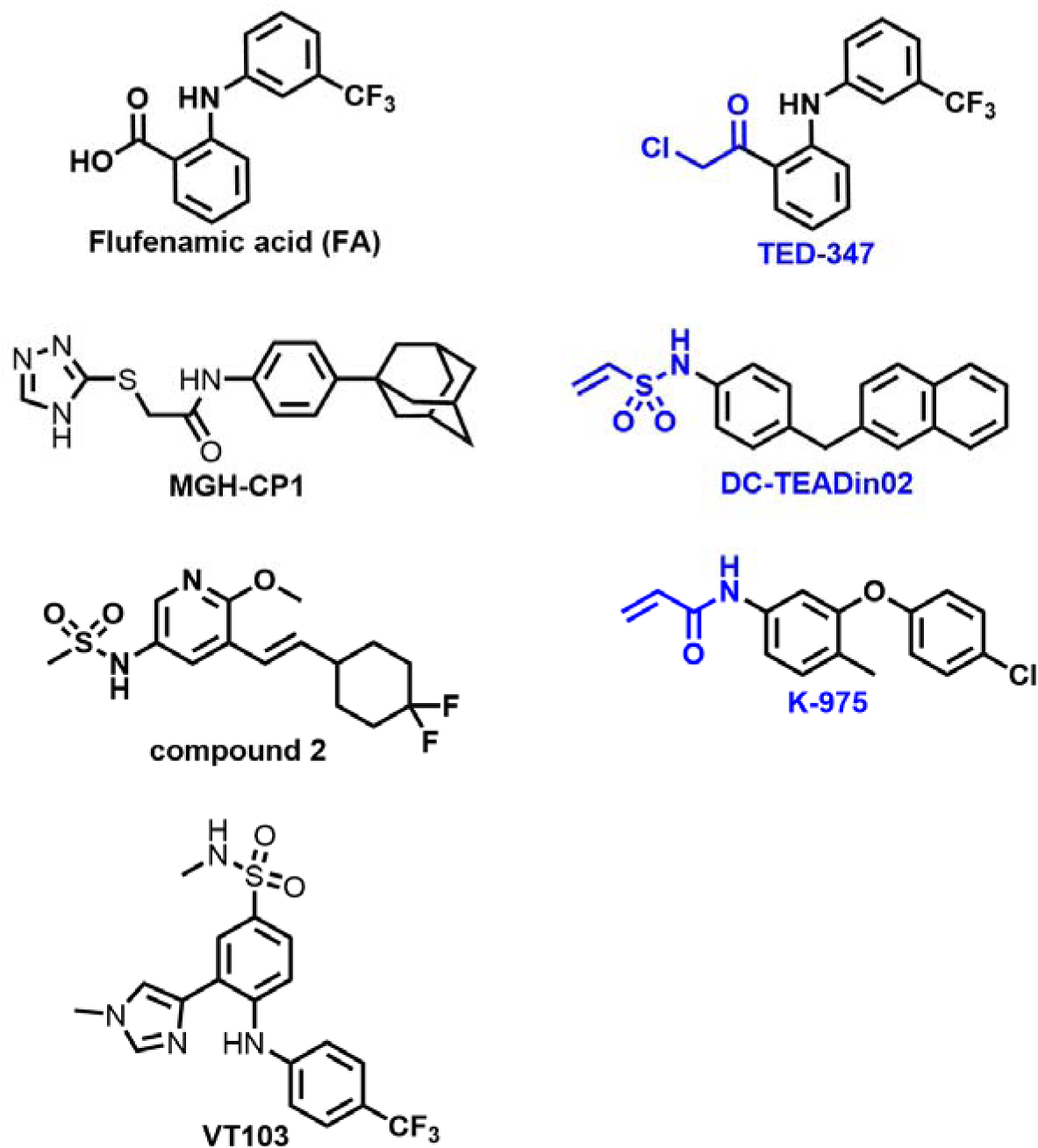
Chemical structures of known TEAD palmitate pocket binders. Covalent binders are labeled in blue.

**Figure S2.**
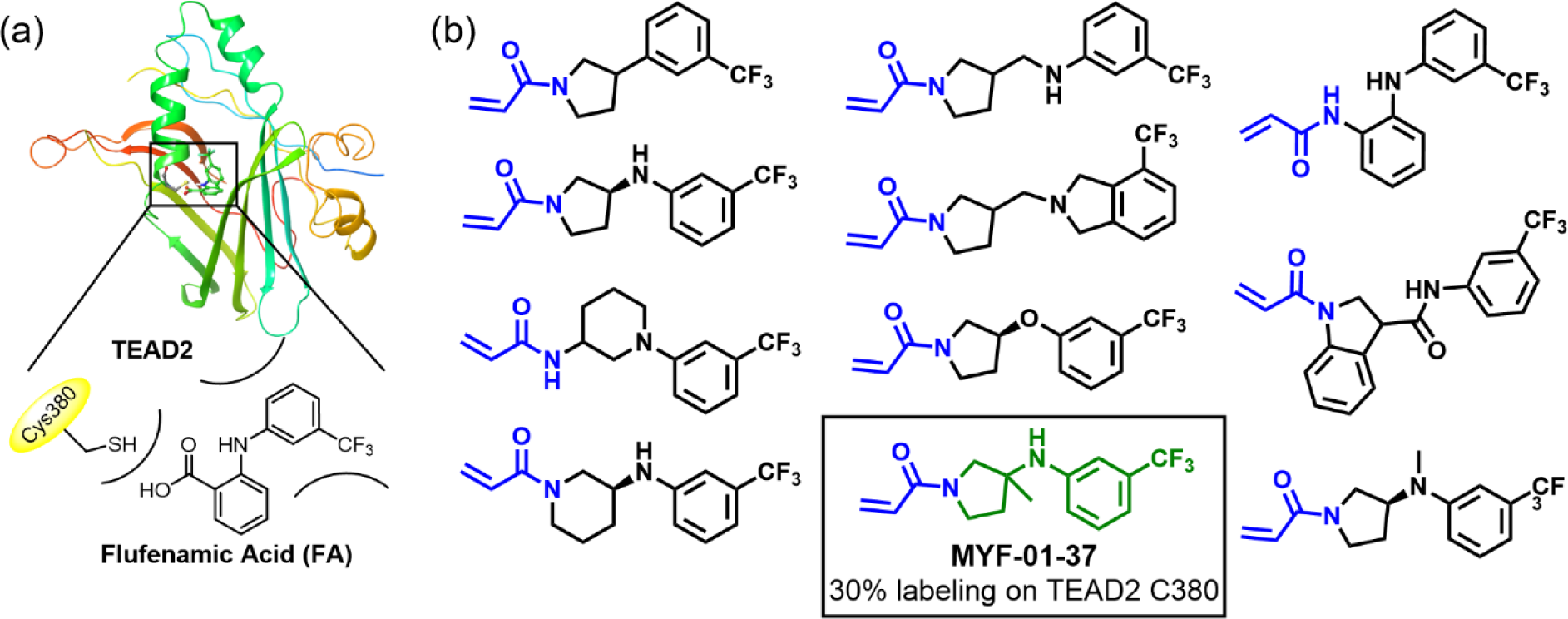
Rationale of covalent fragments design. (**a**) Binding model of FA in TEAD2 palmitate pocket. (**b**) Chemical structures of representative covalent fragments. This figure is related to Figure 1a.

**Figure S3.**
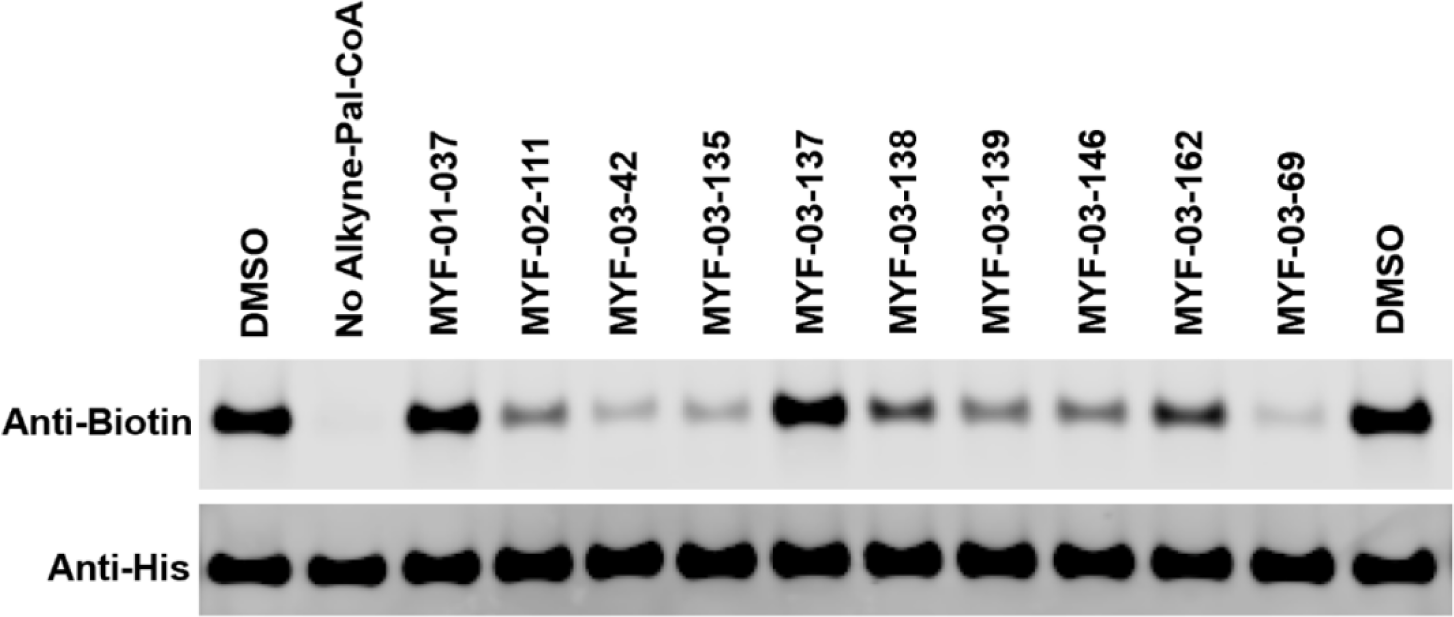
Original western blot image of palmitoylation assay in YBD protein of TEAD2. This figure is related to Figure 1d-e.

**Figure S4.**
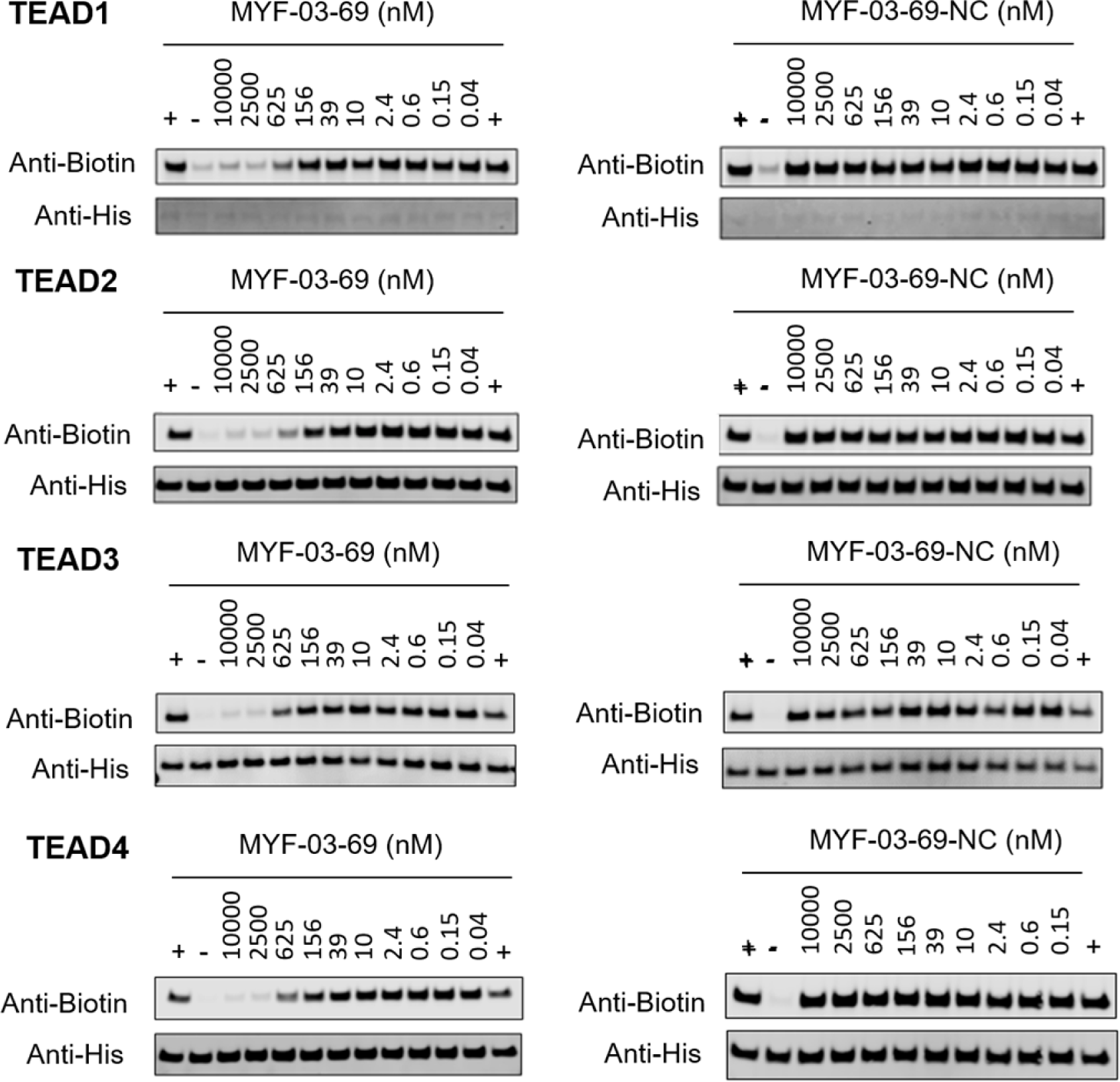
Original western blot images of palmitoylation assays in YBD protein of TEAD1-4. This figure is related to Figure 2e.

**Figure S5.**
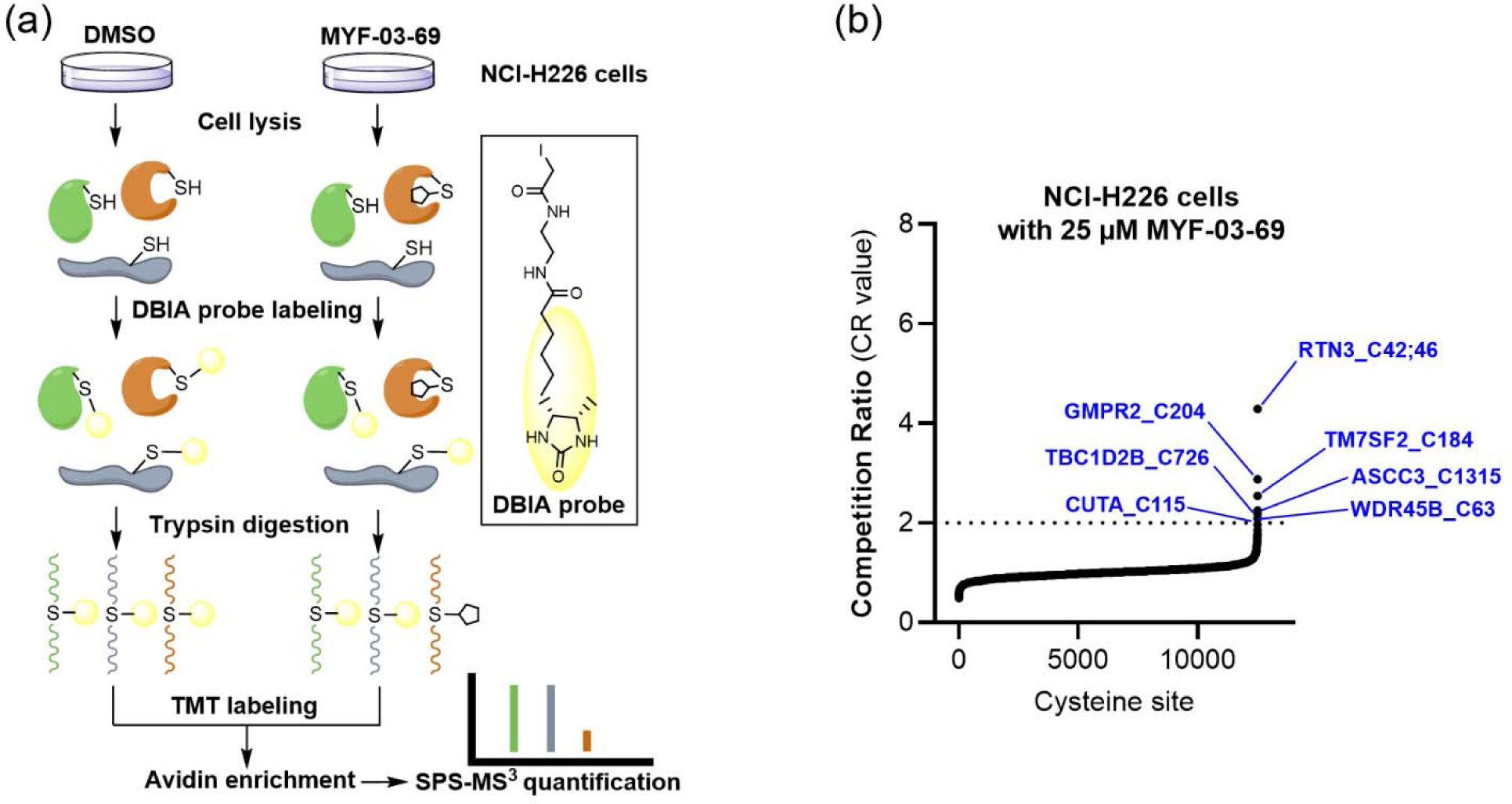
MYF-03-69 exhibited low reactivity towards the cysteine proteome. (**a**) Overview of the SLC-ABPP proteome-wide selectivity approach for profiling of MYF-03-69. NCI-H226 cells were treated with 0.5, 2, 10 or 25 μM MYF-03-69 and DMSO over 3 hours in triplicates before cell lysis followed by labeling with DBIA probe. The SLC-ABPP approach was used to profile competition of MYF-03-69 in proteome cysteine labeling. Competition ratio (CR) was calculated in order to quantitatively assess labeling at every cysteine residue. (**b**) The 7 cysteine sites that were significantly labeled (i.e. exhibited >50% conjugation or CR>2) by 25 µM of MYF-03-69 are in blue, all of which exhibited dose-dependent engagement.

**Figure S6.**
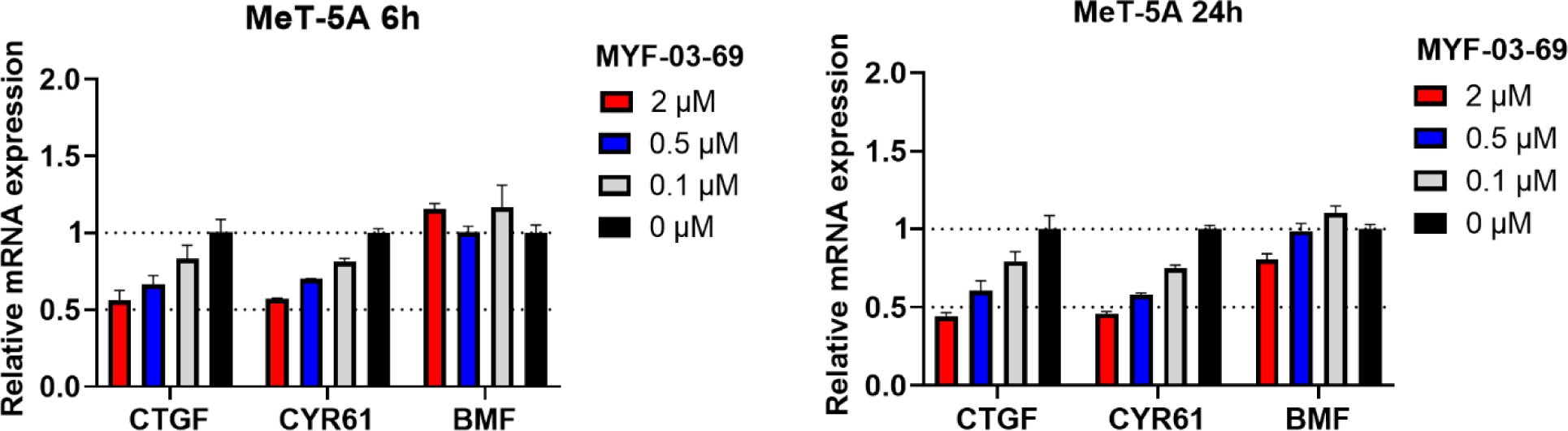
MYF-03-69 showed comparatively weak inhibition on YAP-TEAD downstream gene expression in normal mesothelium cell MeT-5A upon 6-hour and 24-hour treatment. This figure is related to Figure 4b-c.

**Figure S7.**
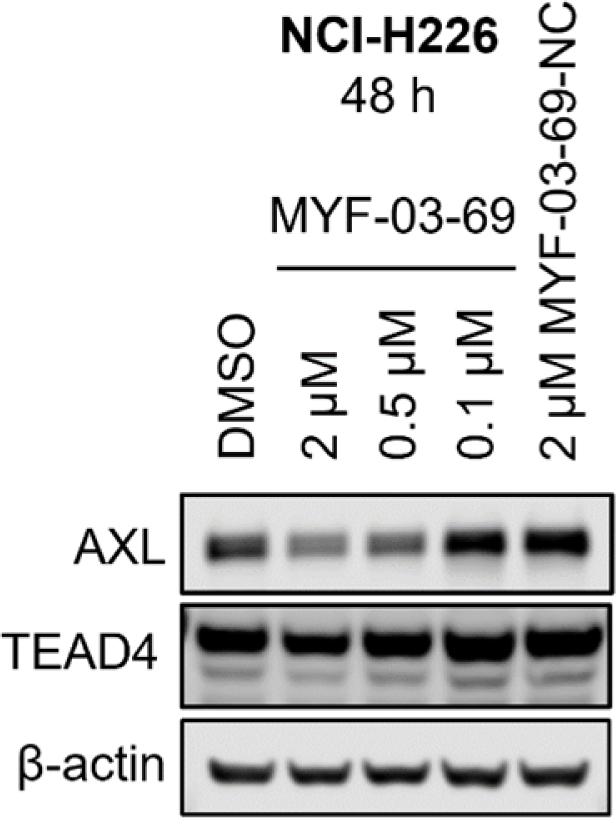
MYF-03-69 downregulated product of canonical YAP downstream gene *AXL* with minimal effect on TEAD stability. This figure is related to Figure 4d.

**Figure S8.**
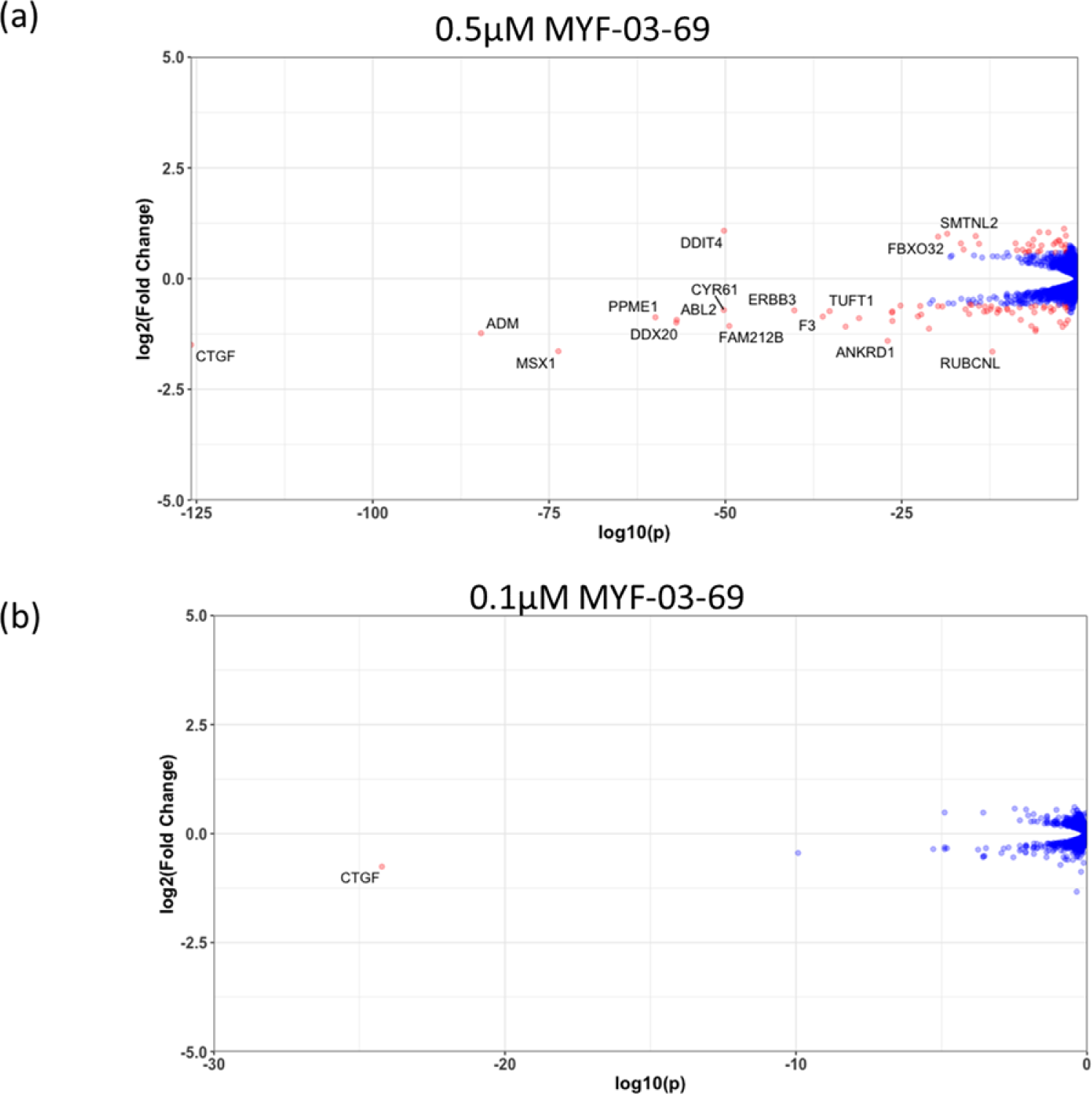
The number of genes with significantly altered expression level decreased with lower concentration treatment in NCI-H226 cells. (**a**) 0.5 μM MYF-03-69, (**b**) 0.1 μM MYF-03-69. This figure is related to Figure 4e-f. Details are in Supplementary Dataset 2.

**Figure S9.**
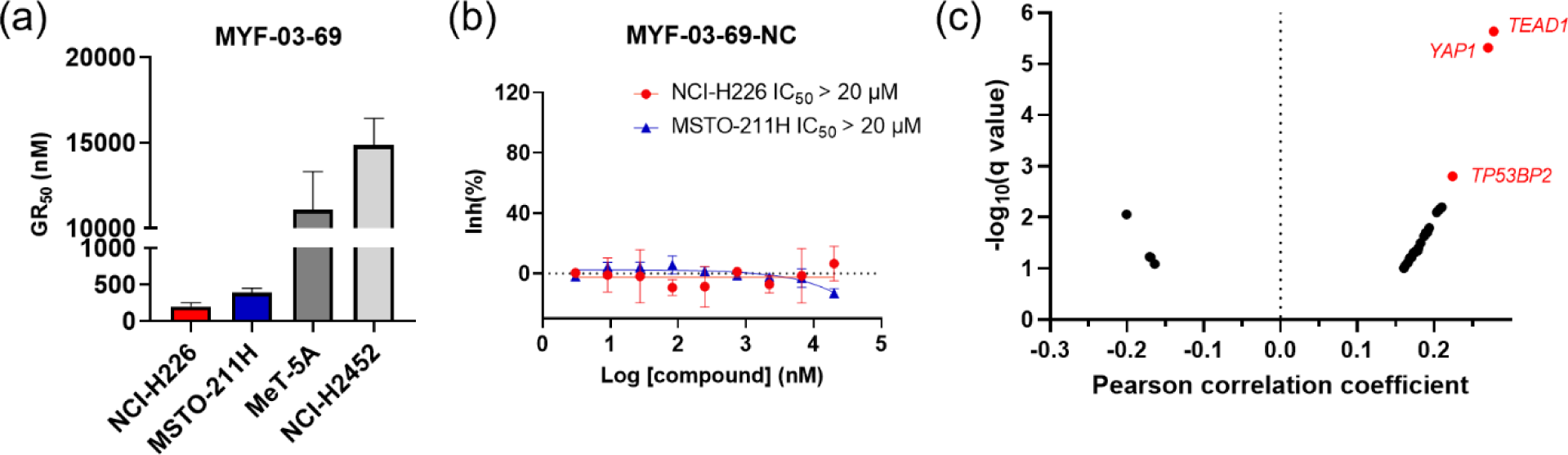
(**a**) The calculated GR_50_ values of MYF-03-69 in 5-day proliferation assay on mesothelioma cells. NCI-H226 and MSTO-211H are Hippo signaling defective mesothelioma cells. NCI-H2452 is Hippo signaling intact mesothelioma cell. MeT-5A is non-cancerous mesothelium cell. This figure is related to Figure 5a. (**b)** MYF-03-69-NC did not inhibit cell proliferation of NCI-H226 and MSTO-211H cells in 5-day treatment. This figure is related to Figure 5a. (**c**) Correlation analysis between compound PRISM sensitivity (log2.AUC of each cell line) and dependency of certain gene (CRISPR knockout score for each cell line, from DepMap Public 20Q4 Achilles_gene_effect.csv dataset) across the PRISM cell line panel. The Pearson correlation coefficients (X axis) and associated p-values were computed. Positive correlations correspond to dependency correlating with increased sensitivity. The q-values (a corrected significance value accounting for false discovery rate) are computed from p-values using the Benjamini Hochberg algorithm. Associations with q-values above 0.1 are filtered out. Top 3 correlated genes are in red. Details are in **Supplementary Dataset 4**.

**Figure S10.**
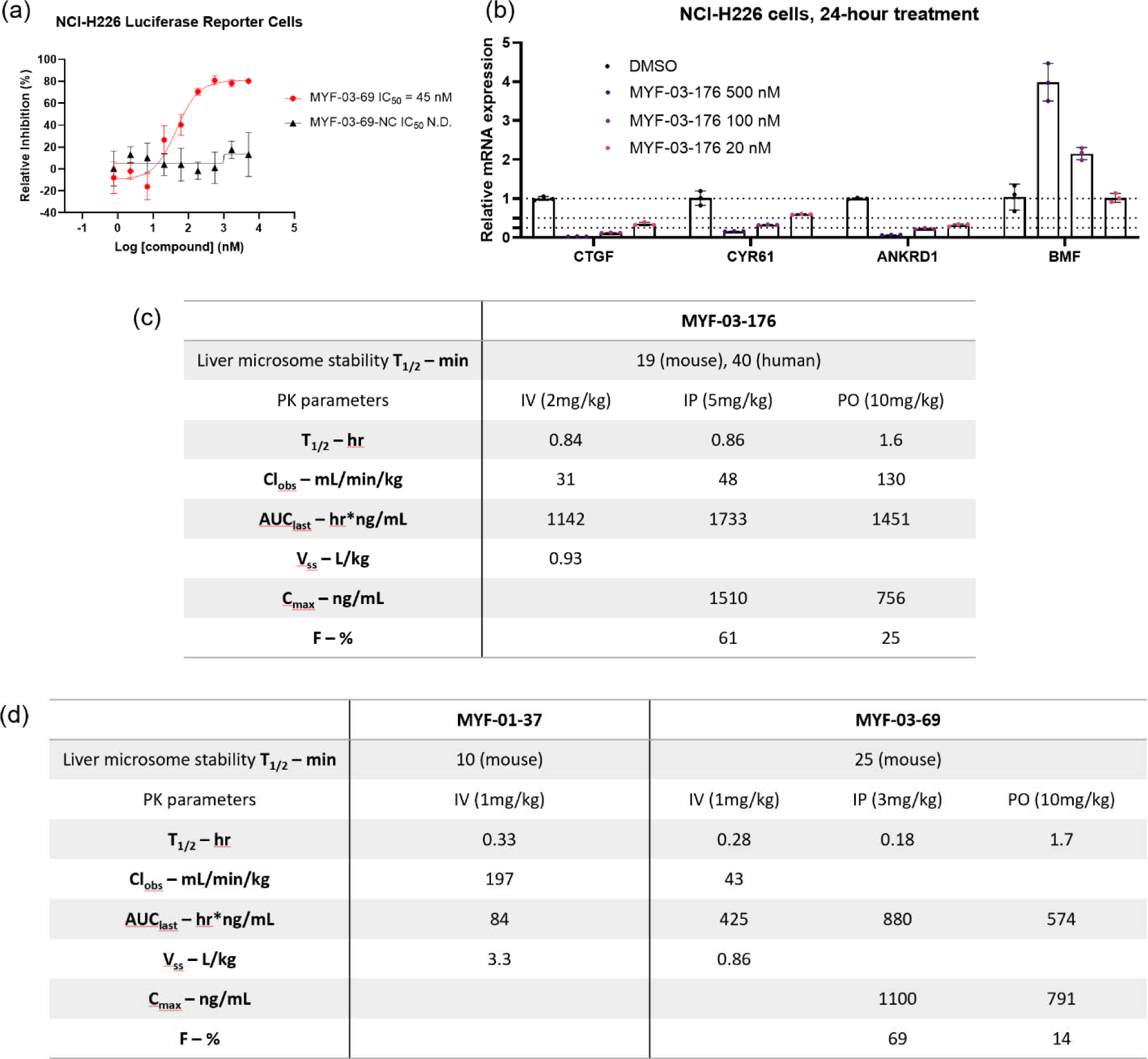
(**a**) MYF-03-69, but not MYF-03-69-NC, inhibits YAP-TEAD transcriptional activity in NCI-H226 luciferase reporter cells. This figure is related to Figure 6b-c. (**b**) MYF-03-176 downregulates YAP target genes and upregulates a pro-apoptotic gene *BMF*. (**c**-**d**) Liver microsome stability and PK parameters comparison of MYF-03-176, MYF-03-69 and MYF-01-37.

